# Gut microbial phenylalanine metabolism contributes to MASLD in a dietary protein-dependent manner

**DOI:** 10.64898/2026.09.28.754936

**Authors:** Yi Rou Bah, Dayang Nurul Asyiqin Binte Mustafa, Frisma Eri Saputri, Takuo Emoto, Sunny H Wong, Rinkoo Dalan, Wei-Kai Wu, Nguan Soon Tan, Torsten Wuestefeld, Kazuyuki Kasahara

## Abstract

Metabolic dysfunction-associated steatotic liver disease (MASLD) affects nearly a third of adults worldwide, yet how diet shapes the microbiota’s contribution remains unclear. In a diet-induced mouse model varying protein content independently of caloric intake, microbiota depletion ameliorated hepatic steatosis and liver injury under protein-sufficient but not protein-restricted conditions. Caecal and plasma metabolomics implicated microbial phenylalanine catabolism and its host conjugates phenylacetylglutamine (PAGln) and phenylacetylglycine (PAGly). PAGln supplementation exacerbated steatosis and liver injury without altering body weight, adiposity or glucose homeostasis. Individual-antibiotic perturbation with metagenomics linked bacterial phenylalanine catabolic capacity to disease severity, and in gnotobiotic mice, genetic disruption of bacterial phenylalanine-to-phenylacetic acid conversion lowered circulating conjugates and attenuated steatosis and fibrosis. In a human cohort, circulating PAGln was selectively elevated in cardiometabolic MASLD. These findings identify dietary protein availability as a modifier of the contribution of microbial phenylalanine metabolism to MASLD and highlight PAGln as a microbiota-derived metabolite that exacerbates hepatic disease.

## Introduction

Metabolic dysfunction-associated steatotic liver disease (MASLD) is the most prevalent chronic liver disease worldwide, affecting nearly one-third of the adult population.^1,2^ It encompasses simple steatosis and its more severe inflammatory form, metabolic dysfunction-associated steatohepatitis (MASH), which may progress to fibrosis, cirrhosis and hepatocellular carcinoma (HCC).^3^ Steatosis may remain clinically stable in some individuals, but a subset progresses to MASH and fibrosis, substantially increasing the risk of liver-related complications and mortality.^3,4^ MASLD is closely intertwined with systemic cardiometabolic dysfunction, commonly coexisting with obesity, insulin resistance, hypertension, dyslipidaemia and type 2 diabetes.^5^ These metabolic comorbidities are important determinants of disease progression, with obesity and type 2 diabetes among the strongest risk factors for advanced fibrosis, cirrhosis and HCC.^6^ Among the multifactorial triggers shaping its progression, dietary exposure represents a major modifiable risk factor.

Dietary protein content has emerged as an important yet understudied modulator of metabolic health, with both its source and quality implicated in metabolic disease in rodents and humans^7–10^. In the Rotterdam Study, higher animal protein intake was associated with increased odds of MASLD among overweight older adults.^11^ Similarly, among patients with biopsy-proven steatotic liver disease, protein intake above 17.3% of total energy was independently associated with a fivefold increase in the odds of steatohepatitis.^12^ Dietary amino acids have also been shown to contribute to hepatic lipogenesis.^13^ However, the mechanisms linking dietary protein intake to MASLD, particularly the extent to which these effects are mediated by the gut microbiota, remain poorly understood.

The gut microbiota is increasingly recognised as a mediator of these diet-host interactions in MASLD, as undigested dietary protein reaching the colon can undergo proteolytic fermentation by gut bacteria, generating an array of bioactive metabolites including branched-chain fatty acids, aromatic amino acid derivatives, short-chain amines, hydrogen sulphide and ammonia^14^. These metabolites enter the portal circulation and can affect hepatic metabolism, depending on their chemical identity and physiological context^15,16^. For example, phenylalanine can be converted by the microbiota to phenylpyruvic acid (PPY) and subsequently phenylacetic acid (PAA) through distinct microbial enzymatic pathways. PAA is then conjugated by the host to phenylacetylglutamine (PAGln) or phenylacetylglycine (PAGly), metabolites that have been associated with cardiometabolic diseases, including coronary heart disease, heart failure and chronic kidney disease^17,18,19^. However, whether this pathway causally contributes to MASLD, and whether its relevance depends on dietary protein availability, remains unresolved.

Here, we show that dietary protein availability modulates the contribution of the gut microbiota to MASLD, identify microbial phenylalanine metabolism as a candidate pathway linking dietary protein, the gut microbiota, and MASLD, and provide genetic evidence that a defined bacterial phenylalanine catabolic function contributes to MASLD development.

## Results

### Dietary protein availability modifies the hepatic response to gut microbiota depletion

Previous studies have demonstrated a role for the gut microbiota in diet-induced MASLD models.^20–24^ We therefore first assessed the effect of microbiota depletion in the Liver Disease Progression Aggravation Diet (LIDPAD) model,^25^ which combines a high-fat, high-cholesterol diet with thermoneutral housing. Microbiota depletion with an antibiotic mixture (ampicillin, vancomycin, metronidazole, neomycin) reduced faecal bacterial DNA content by approximately four orders of magnitude and attenuated body weight gain, adiposity and glucose intolerance, but unexpectedly did not improve any histological measure of MASLD severity (**Supplementary Figure 1A-I**). This dissociation prompted us to investigate whether the dietary context modifies the contribution of the gut microbiota to hepatic disease.

Given the relatively low protein content of the LIDPAD diet (∼11% of total energy), we next asked whether dietary protein availability influences the hepatic response to microbiota depletion. Mice were fed isocaloric LIDPAD diets containing either 21% (LID21) or 7% (LID7) of total energy from protein, with or without antibiotic-mediated microbiota depletion, for 12 weeks under thermoneutral housing (**Figure 1A**). Antibiotic treatment attenuated body weight gain in both dietary groups throughout the study (**Figure 1B**). At endpoint, antibiotic treatment reduced body weight and liver-to-body weight ratio in LID21-fed mice but not in LID7-fed mice, whereas it reduced fat-to-body weight ratio in both (**Figure 1C**). Both LID21- and LID7-fed mice showed a trend towards improved glucose and insulin tolerance after antibiotic treatment (**Supplementary Figures 2A-B**). Antibiotic treatment reduced proinflammatory Ly6C^hi^ monocytes in LID21-fed mice but not in LID7-fed mice (**Supplementary Figure 3A-C**).

**Figure 1.**
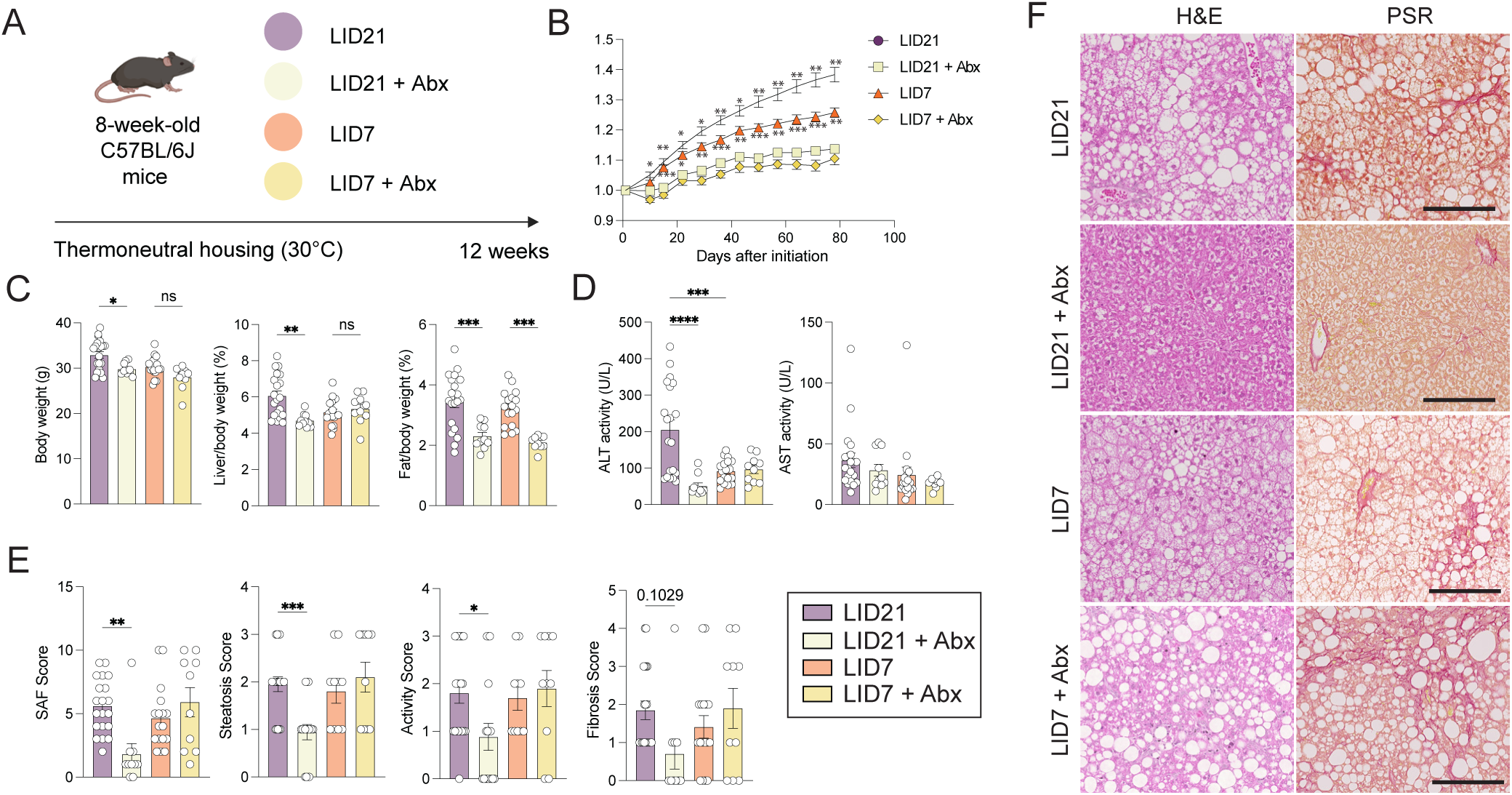
The influence of gut microbiota on MASLD pathogenesis is modulated by dietary protein availability. **(A)** Schematic of the experimental design. Eight-week-old C57BL/6J mice were fed LIDPAD diets providing either 21% (LID21) or 7% (LID7) of total energy from protein, with or without antibiotic treatment, for 12 weeks under thermoneutral housing conditions. **(B)** Body weight normalised to initial body weight over the study period. **(C)** Body weight, liver/body weight ratio, and fat/body weight ratio at the end of the study. **(D)** Plasma ALT and AST activities. **(E)** Composite SAF score, steatosis score, activity score, and fibrosis score. **(F)** Representative histological images of liver sections stained with H&E and PSR. Scale bars, 50 µm. Data are represented as mean ± SEM, n = 20 (LID21), 10 (LID21 + Abx), 18 (LID7), 10 (LID7 + Abx). Statistical significance was determined by one-way ANOVA with Tukey’s post hoc test for (C-D), Kruskal-Wallis with Dunn’s post hoc test for (E) and two-way repeated-measures ANOVA for (B). *p < 0.05, **p < 0.01, ***p < 0.001, ****p < 0.0001; ns, not significant.

The hepatic response to microbiota depletion differed between the two dietary groups. Plasma ALT activity was markedly reduced by antibiotic treatment in LID21-fed mice but unchanged in LID7-fed mice, with AST unaffected in both (**Figure 1D**). Consistently, antibiotic treatment reduced the composite SAF, steatosis and activity scores in LID21-fed mice, with a trend towards reduced fibrosis, whereas none of these scores differed between LID7 and LID7 + Abx mice (**Figure 1E-F**).

Together, these data show that microbiota depletion was associated with marked improvements in hepatic steatosis and liver injury in LID21-fed mice, whereas comparable improvements were not detected in LID7-fed mice. These findings suggest that dietary protein availability may modulate the contribution of the gut microbiota to MASLD progression.

### Metabolomics identifies phenylalanine metabolism as a candidate pathway linking dietary protein, the gut microbiota, and MASLD

To identify metabolic pathways potentially linking dietary protein, the gut microbiota, and MASLD, we performed untargeted metabolomics of caecal contents and targeted metabolomics of plasma collected from LID21, LID21 + Abx, LID7, and LID7 + Abx mice (**Figure 2A**). Principal component analysis of the caecal metabolome showed clear separation of antibiotic-treated and untreated mice along PC1 (45% of variance), whereas LID21- and LID7-fed mice were primarily separated along PC2 (13.1% of variance, **Figure 2B**). These findings indicate that microbiota depletion was the dominant driver of caecal metabolome variation, with dietary protein content providing an additional source of variation.

**Figure 2.**
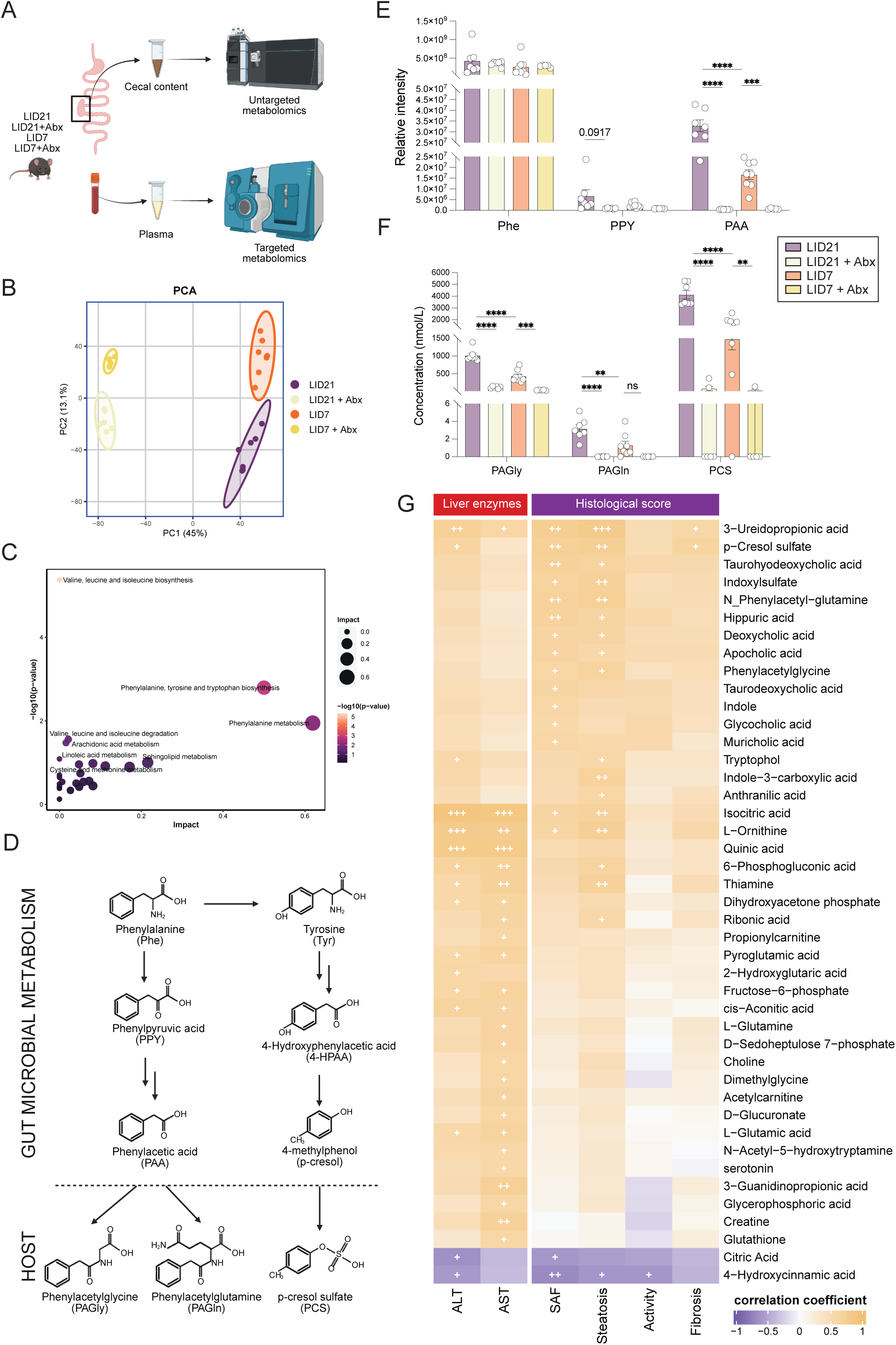
Metabolomics implicates microbial aromatic amino acid metabolism in the dietary protein-microbiota interaction in MASLD. **(A)** Schematic of the metabolomics workflow. Caecal contents were analysed by untargeted metabolomics and plasma samples by targeted metabolomics in LID21, LID21 + Abx, LID7, and LID7 + Abx mice. **(B)** Principal component analysis (PCA) of caecal untargeted metabolomics data across the four groups. **(C)** Pathway enrichment analysis of the 765 caecal metabolites assigned to cluster 7 by k-means clustering. Bubble size represents pathway impact, and colour represents statistical significance (−log10 p value). **(D)** Schematic of the microbial phenylalanine and tyrosine catabolic pathways and subsequent host conjugation, showing conversion of phenylalanine (Phe) to phenylpyruvic acid (PPY) and phenylacetic acid (PAA), and of tyrosine (Tyr) to 4-hydroxyphenylacetic acid (4-HPAA) and p-cresol, with subsequent host conjugation to phenylacetylglycine (PAGly), phenylacetylglutamine (PAGln), and p-cresol sulfate (PCS). **(E)** Relative intensities of caecal Phe, PPY, and PAA measured by untargeted metabolomics. **(F)** Plasma concentrations of PAGly, PAGln, and PCS measured by targeted metabolomics. **(G)** Heatmap of Spearman correlations between plasma metabolites and liver enzyme activities (ALT, AST) or histological scores (SAF, steatosis, activity, and fibrosis). Colour indicates the correlation coefficient. Data are represented as mean ± SEM. n = 7 (LID21), 6 (LID21 + Abx), 8 (LID7), 5 (LID7 + Abx). Statistical significance was determined by one-way ANOVA with Tukey’s post hoc test. *p < 0.05, **p < 0.01, ***p < 0.001, ****p < 0.0001; ns, not significant. For (G), +p < 0.05, ++p < 0.01, +++p < 0.001.

To explore metabolite patterns associated with the phenotypic response across dietary and microbiota-depletion conditions, we first applied k-means clustering to annotated caecal metabolites based on their abundance profiles across the four experimental groups (**Supplementary Figure 4A**). Clusters 4 and 7, comprising 1,066 and 765 metabolites, respectively, showed patterns resembling the phenotypic response and were therefore examined in this exploratory analysis. Pathway analysis of cluster 7 highlighted phenylalanine metabolism, together with phenylalanine, tyrosine and tryptophan biosynthesis and branched-chain amino acid pathways (**Figure 2C**, **Supplementary Figure 4B**).

Because k-means clustering depends on the prespecified number of clusters and subsequent cluster selection, we next tested the robustness of this observation using analyses that did not rely on metabolite clustering. Microbiota depletion broadly altered the caecal metabolome, affecting 5,387 of 7,439 metabolites (72.4%; FDR < 0.05), limiting discrimination at the pathway level: competitive testing identified only secondary bile acid biosynthesis, whereas self-contained testing identified 109 of 138 KEGG pathways (**Supplementary Figure 4C**). We therefore ranked all detected metabolites according to the effect of microbiota depletion. Phenylacetic acid (PAA) ranked within the most strongly depleted 1.2% of the metabolome, and 2-, 3-, and 4-hydroxyphenylacetic acids within the most depleted 4%, whereas phenylalanine and tyrosine themselves were largely unchanged (51st and 54th percentiles, respectively; **Supplementary Figure 4D-F**). We next directly tested whether PAA exhibited a protein-by-microbiota interaction. Protein restriction reduced caecal PAA in microbiota-intact mice (t = −4.24, FDR = 0.002) but not following microbiota depletion (t = 1.87), resulting in a significant protein × antibiotic interaction (t = −4.18, FDR = 0.009; **Supplementary Figure 4G**). Together, these cluster-independent analyses supported a microbiota- and protein-dependent pattern of PAA abundance and preferential depletion of microbial aromatic amino acid catabolites rather than their amino acid substrates.

Gut bacteria can metabolise phenylalanine into phenylpyruvic acid (PPY) and subsequently PAA, which can be conjugated by the host to form phenylacetylglutamine (PAGln) and phenylacetylglycine (PAGly). Similarly, microbial metabolism of tyrosine generates 4-hydroxyphenylacetic acid and p-cresol, with the latter undergoing host conjugation to form p-cresol sulfate (PCS) (**Figure 2D**). Caecal phenylalanine concentration did not differ among groups, whereas PAA was significantly reduced by antibiotic treatment in both LID21- and LID7-fed mice, with a similar trend for PPY (**Figure 2E**). Consistent with these findings, 367 metabolites met the predefined criteria for a microbiota-dependent protein response (**Supplementary Table 1**), including PAA and PCS. Given its strong microbiota dependence, significant protein × antibiotic interaction, and its role as the direct precursor of PAGln and PAGly, we prioritised PAA for further investigation. Notably, PAGln has been implicated in heart failure and cellular senescence.^26,27^

We next quantified the corresponding host-conjugated microbial metabolites in plasma. PAGly, PAGln, and PCS were significantly higher in LID21 than in LID21 + Abx mice and were also higher in LID21 than in LID7 mice. PAGly and PCS were also reduced by antibiotic treatment in LID7-fed mice (**Figure 2F**). Consistent with previous reports that PAGly is the dominant conjugate in mice, whereas PAGln predominates in humans,^17,19^ PAGln concentrations were substantially lower than PAGly across all groups.

We further examined the correlation between circulating metabolites and indices of liver injury and histological disease severity. Phenylalanine- and tyrosine-derived microbial metabolites, together with several secondary bile acids and tryptophan-derived indoles, correlated positively with ALT, AST, SAF, and steatosis scores, while correlations with activity and fibrosis scores were comparatively weak (**Figure 2G**). In contrast, citric acid and 4-hydroxycinnamic acid were negatively correlated with ALT and SAF scores. Together, these complementary metabolomic analyses identified microbial aromatic amino acid metabolism, and phenylalanine-to-PAA metabolism in particular, as a candidate pathway linking dietary protein, the gut microbiota, and MASLD.

### PAGln exacerbates liver steatosis and liver injury in vivo

Given that PAGln is the predominant PAA conjugate in humans and was elevated in LID21-fed, microbiota-intact mice, we first asked whether PAGln directly promotes lipid accumulation in hepatocytes. Treatment of cultured hepatocytes with PAGln did not increase intracellular triglyceride content across the concentrations tested, either alone or in the presence of the steatogenic stimulus OAPA (**Supplementary Figure 5**). PAGly similarly had no effect, whereas PCS produced a modest dose-dependent increase in triglyceride accumulation under basal conditions. These findings suggested that PAGln does not directly promote hepatocyte triglyceride accumulation.

We therefore next asked whether PAGln could exacerbate MASLD in vivo. Eight-week-old B6 mice fed the LIDPAD diet were administered vehicle or PAGln for 12 weeks under thermoneutral housing conditions (**Figure 3A**). PAGln treatment did not significantly alter relative weight gain over the treatment period, or final body weight, liver-to-body weight ratio, or fat-to-body weight ratio (**Figures 3B-C**). PAGln treatment also did not significantly affect glucose or insulin tolerance (**Supplementary Figure 6A-B**).

**Figure 3.**
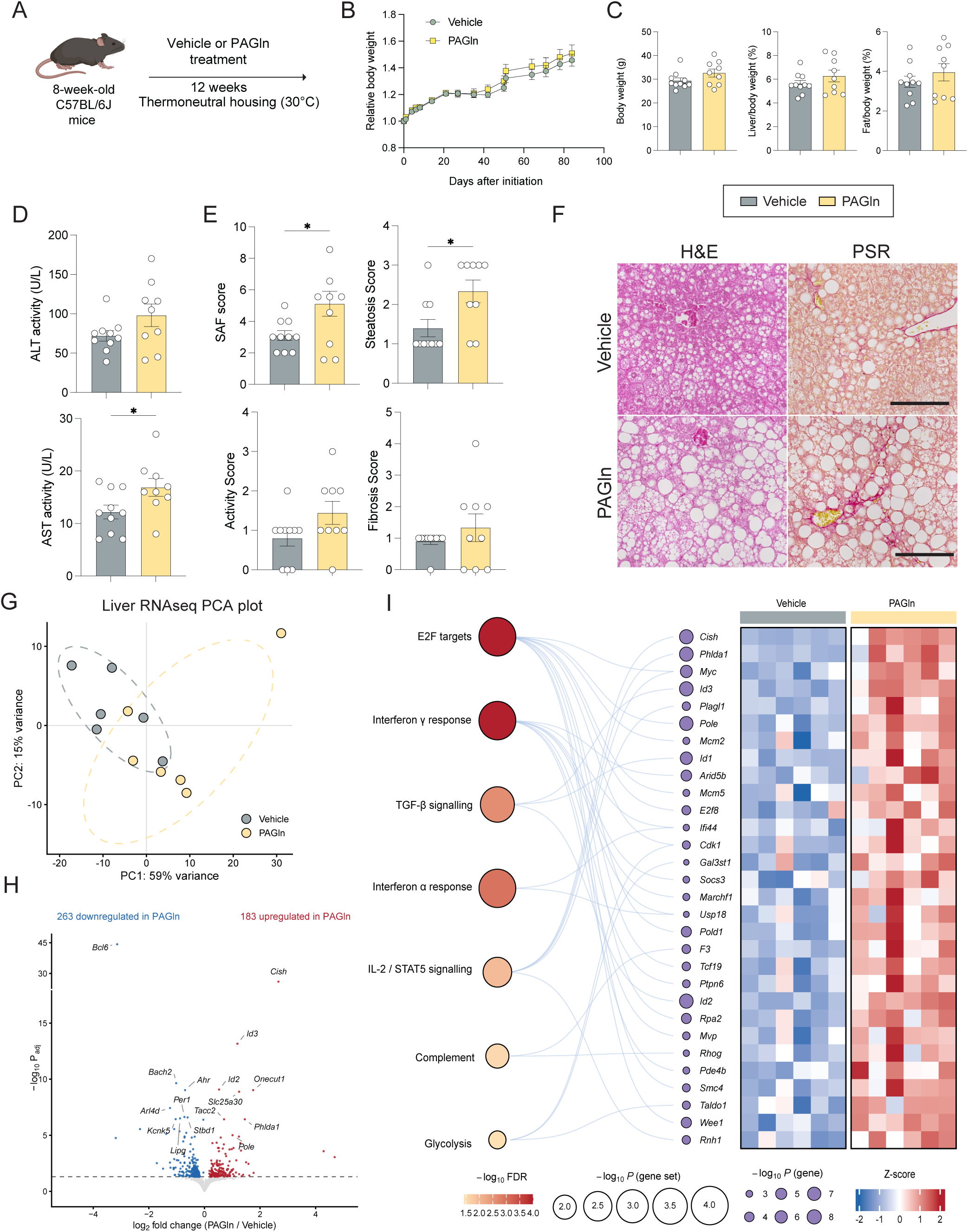
PAGln administration exacerbates diet-induced MASLD. **(A)** Schematic of the experimental design. Eight-week-old C57BL/6J mice were fed a LIDPAD diet and administered vehicle or phenylacetylglutamine (PAGln) for 12 weeks under thermoneutral housing conditions. **(B)** Body weight normalised to initial body weight over the study period. **(C)** Body weight, liver/body weight ratio, and fat/body weight ratio at the end of the study. **(D)** Plasma ALT and AST activities. **(E)** Composite SAF score, steatosis score, activity score, and fibrosis score. **(F)** Representative histological images of liver sections stained with haematoxylin and eosin (H&E) and picrosirius red (PSR). Scale bars, 50 µm. **(G)** Principal component analysis of liver RNA-seq data from vehicle- and PAGln-treated mice using variance-stabilised counts. Each point represents one animal; dashed ellipses indicate 95% confidence regions. Percentages indicate the variance explained by each principal component. **(H)** Volcano plot of differential gene expression in PAGln-versus vehicle-treated livers. Red and blue points indicate significantly upregulated and downregulated genes, respectively (adjusted *P* < 0.05); selected genes are labelled. **(I)** Gene set enrichment analysis against the mouse Hallmark gene set collection. Left, significantly positively enriched Hallmark gene sets; centre, selected leading-edge genes contributing to the enriched gene sets; right, heatmap of z-scored variance-stabilised expression of the same genes. Data are represented as mean ± SEM. For (B-F), n = 10 (vehicle), 9 (PAGln). For (G-I), n = 6 per group. Statistical significance was determined by unpaired two-tailed t-test for (C-D) and by Mann-Whitney test for (E); for (G-I), differential expression was assessed in DESeq2 by Wald test with Benjamini-Hochberg correction, and gene set enrichment was performed using fgsea with Benjamini–Hochberg correction. *p < 0.05.

Plasma AST activity was significantly increased by PAGln treatment, whereas ALT activity showed a non-significant trend toward an increase (**Figure 3D**). Consistent with increased liver injury, PAGln-treated mice had significantly higher composite SAF and steatosis scores than vehicle-treated mice, while activity and fibrosis scores were not significantly different (**Figure 3E-F**). Thus, PAGln exacerbated hepatic steatosis and liver injury without detectable changes in body weight, adiposity, or systemic glucose homeostasis.

We next performed bulk RNA sequencing of liver tissues to characterise the hepatic transcriptional changes associated with PAGln treatment. Principal component analysis showed partial separation between vehicle- and PAGln-treated mice along PC1, which accounted for 59% of the variance (**Figure 3G**). Differential expression analysis identified 446 differentially expressed genes, with 183 upregulated and 263 downregulated in PAGln-treated livers (**Figure 3H**). Gene set enrichment analysis identified seven positively enriched Hallmark gene sets, including E2F targets, interferon α and γ responses, TGF-β signalling, IL-2/STAT5 signalling, complement and glycolysis (**Figure 3I**). These changes were driven by genes involved in cell-cycle regulation, interferon responses and cytokine signalling, whereas no lipid metabolism-related Hallmark gene set reached significance. Together, these findings show that PAGln treatment is associated with hepatic transcriptional changes involving immune-inflammatory pathways accompanied by stress-response and metabolic adaptation programs.

We next examined whether PCS, another host-conjugated microbial metabolite associated with the LID21-fed, microbiota-intact condition, exerted similar effects. In a separate experiment, mice were fed a LIDPAD diet for 12 weeks and received either water alone or 0.05% PCS in their drinking water (**Supplementary Figure 7A**). PCS supplementation showed a trend toward increased body weight without significantly affecting epididymal fat-to-body weight or liver-to-body weight ratios (**Supplementary Figure 7B-C**). Glucose and insulin tolerance were also not significantly affected (**Supplementary Figure 7D-E**). Unlike PAGln, PCS supplementation reduced plasma AST activity, with a non-significant trend toward reduced ALT activity (**Supplementary Figure 7F**). Histological assessment showed an increased activity score in PCS-treated mice, whereas SAF, steatosis, and fibrosis scores were not changed (**Supplementary Figure 7G-H**).

Taken together, these findings show that PAGln exacerbates hepatic steatosis and liver injury in diet-induced MASLD without altering body weight, adiposity, or systemic glucose homeostasis and is accompanied by hepatic transcriptional changes involving immune-inflammatory pathways and metabolic remodelling. PCS did not reproduce the PAGln-associated phenotype, indicating distinct hepatic effects of these host-conjugated microbial metabolites.

### Individual antibiotics identify gut bacterial taxa associated with phenylalanine catabolism and MASLD severity

To identify specific gut microbiota components associated with MASLD severity, 8-week-old mice fed the LID21 diet received ampicillin, vancomycin, metronidazole, or neomycin, or were left untreated, for 12 weeks under thermoneutral housing conditions (**Figure 4A**). Antibiotic treatment differentially affected body weight trajectories, with ampicillin- and metronidazole-treated mice showing the greatest attenuation of body weight (**Figure 4B**). Final body weight did not differ significantly among groups. Ampicillin treatment significantly reduced the liver-to-body weight ratio compared with untreated controls, whereas metronidazole treatment significantly reduced the fat-to-body weight ratio (**Figure 4C**).

**Figure 4.**
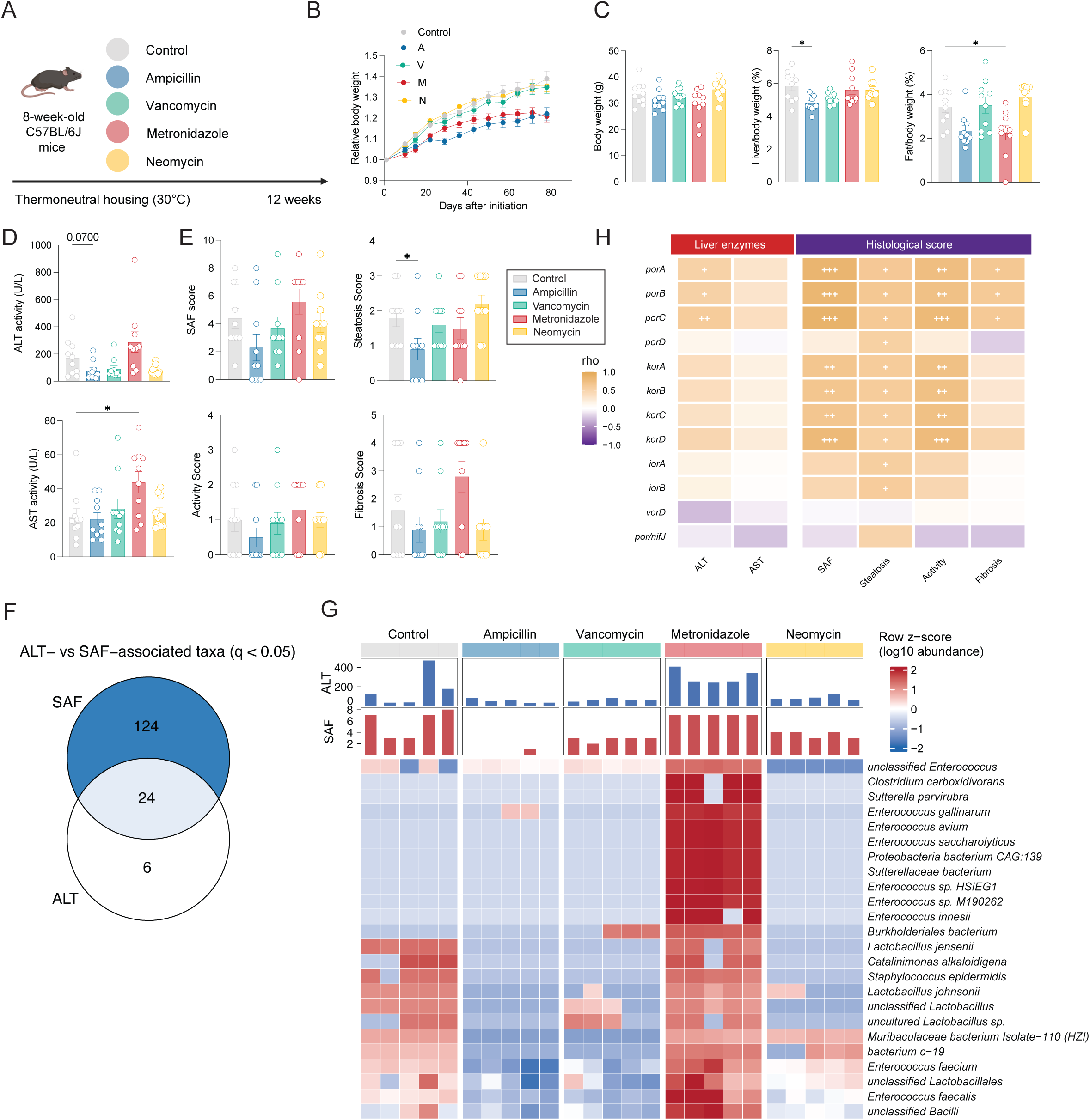
Individual antibiotic perturbation identifies gut bacterial taxa and candidate phenylalanine catabolic genes associated with MASLD severity. **(A)** Schematic of the experimental design. Eight-week-old C57BL/6J mice were fed a LIDPAD diet providing 21% of total energy from protein (LID21) and received no antibiotics (control) or individual treatment with ampicillin, vancomycin, metronidazole, or neomycin for 12 weeks under thermoneutral housing conditions. **(B)** Body weight normalised to initial body weight over the study period. **(C)** Body weight, liver/body weight ratio and fat/body weight ratio at the end of the study. **(D)** Plasma ALT and AST activities. **(E)** Composite SAF score, steatosis score, activity score, and fibrosis score. **(F)** Venn diagram of bacterial taxa significantly associated with plasma ALT activity and/or SAF score (q < 0.05). **(G)** Heatmap showing the relative abundance of the 24 bacterial taxa significantly associated with both ALT and SAF for each mouse. Colour indicates the row z-score of log10-transformed relative abundance. Bar plots above the heatmap show the corresponding ALT activity and SAF score. **(H)** Heatmap of Spearman correlations between the abundance of candidate 2-oxoacid:ferredoxin oxidoreductase genes implicated in aromatic amino acid metabolism (*porA–D*, *korA– D*, *iorA/B*, *vorD*, and *por/nifJ*) and plasma liver enzyme activities (ALT and AST) or histological scores (SAF, steatosis, activity, and fibrosis). Data in (B-E) are represented as mean ± SEM. For (B-E), n = 10 per group. Statistical significance was determined by two-way repeated-measures ANOVA for (B), one-way ANOVA with Tukey’s post hoc test for (C-D), and Kruskal–Wallis test with Dunn’s post hoc test for (E). *p < 0.05. Associations between bacterial taxa and ALT or SAF score were assessed by MaAsLin2 in (F) and (G). Correlations in (H) were assessed using Spearman correlation with Benjamini-Hochberg correction; +p < 0.05, ++p < 0.01, +++p < 0.001.

Plasma ALT activity showed a trend toward reduction with ampicillin treatment, whereas AST activity was significantly higher in metronidazole-treated mice than in untreated controls (**Figure 4D**). Histological assessment showed a significant reduction in steatosis score in ampicillin-treated mice, while composite SAF, activity, and fibrosis scores did not differ significantly across groups (**Figure 4E, Supplementary Figure 8A**). These findings indicate that individual antibiotics differentially affect MASLD-related phenotypes.

Quantification of bacterial 16S rRNA gene copies in faecal samples showed that bacterial load was reduced by all antibiotic treatments, with the greatest reduction observed following ampicillin treatment (**Supplementary Figure 8B**). Both Shannon and Chao1 indices were significantly lower in every antibiotic-treated group than in untreated LID21 controls, with the greatest reduction in richness seen with vancomycin (**Supplementary Figure 8C**). Principal coordinates analysis further demonstrated distinct microbial community structures across antibiotic treatment groups (**Supplementary Figure 8D-E**), accompanied by treatment-specific shifts in taxonomic composition (**Supplementary Figure 8F**).

To identify bacterial taxa associated with liver injury and histological disease severity, we used MaAsLin2 to identify taxa associated with plasma ALT activity and SAF score (q < 0.05). A total of 24 taxa were associated with both ALT and SAF score, whereas 124 and 6 taxa were uniquely associated with SAF score and ALT activity, respectively (**Figure 4F**). Examination of the 24 shared taxa revealed distinct abundance patterns across antibiotic treatment groups. Several taxa, including multiple *Enterococcus* species, *Clostridium carboxidivorans*, and members of *Sutterellaceae* and *Proteobacteria*, were enriched in metronidazole-treated mice, which exhibited relatively high ALT activity and SAF scores (**Figure 4G**).

Given the identification of phenylalanine metabolism in our metabolomic analyses, we next examined microbial genes potentially involved in converting PPY to PAA. The 2-oxoacid:ferredoxin oxidoreductase gene families *por, kor, ior, vor* encode enzymes with the potential to catalyse this reaction. The relative abundances of several of these genes, particularly *porA-C* and *korA-D*, were positively correlated with liver enzymes, SAF, steatosis, activity, and fibrosis scores (**Figure 4H**). We therefore screened the 24 taxa associated with both ALT and SAF score for the presence of these genes. Several of the metronidazole-enriched *Enterococcus* species carried *por*/*nifJ* homologues, while *Muribaculaceae bacterium Isolate-110_HZI* and *bacterium c-19* encoded a broader repertoire of candidate genes involved in 2-oxoacid metabolism (**Supplementary Figure 8G**). Together, these findings identify bacterial taxa associated with MASLD severity and implicate phenylpyruvate/indolepyruvate:ferredoxin oxidoreductases as candidate contributors to microbial PAA production.

### Bacterial BT0430 mediates aromatic amino acid metabolism and contributes to MASLD in a gnotobiotic model

The preceding analyses implicated phenylpyruvate/indolepyruvate:ferredoxin oxidoreductases as candidate contributors to microbial PAA production. o test whether a representative phenylpyruvate:ferredoxin oxidoreductase can functionally contribute to PAA production and MASLD, we focused on BT0430 in *Bacteroides thetaiotaomicron* as a genetically tractable representative of this enzyme family (**Figure 5A**). A markerless ΔBT0430 deletion strain was generated by allelic exchange (**Supplementary Figure 9A-B**) and exhibited growth comparable to that of wild-type *B. thetaiotaomicron* in vitro (**Figure 5B**). When wild-type or ΔBT0430 bacteria were cultured in medium supplemented with phenylalanine, PAA was detected only in cultures of the wild-type strain, demonstrating that BT0430 is required for PAA production by *B. thetaiotaomicron* under these conditions (**Figure 5C**).

**Figure 5.**
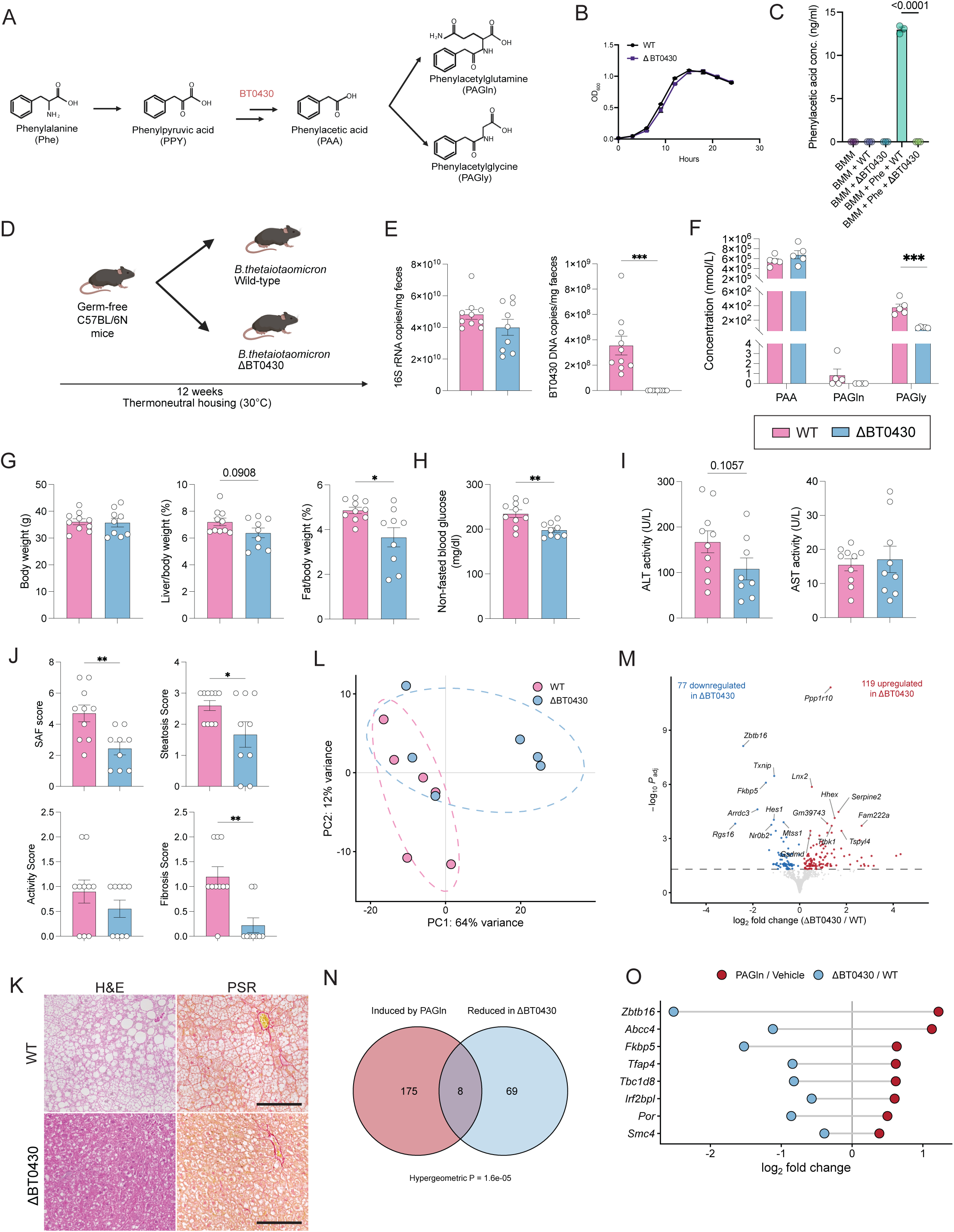
Disruption of a bacterial phenylalanine-catabolic gene attenuates MASLD in gnotobiotic mice. **(A)** Schematic of phenylalanine metabolism, highlighting the proposed role of *Bacteroides thetaiotaomicron* BT0430 in the conversion of phenylpyruvic acid (PPY) to phenylacetic acid (PAA), followed by host conjugation to phenylacetylglutamine (PAGln) and phenylacetylglycine (PAGly). **(B)** Growth curves (OD600) of wild-type (WT) and ΔBT0430 *B. thetaiotaomicron* over 24 h in culture. **(C)** PAA concentrations in Bacteroides minimal medium (BMM) following culture of WT or ΔBT0430 *B. thetaiotaomicron*, with or without phenylalanine (Phe) supplementation. **(D)** Schematic of the experimental design. Germ-free C57BL/6N mice were monocolonised with WT or ΔBT0430 *B. thetaiotaomicron* and fed a LIDPAD diet providing 21% of total energy from protein for 12 weeks under thermoneutral housing conditions. **(E)** Faecal 16S rRNA gene and BT0430 gene copy numbers in WT- and ΔBT0430-colonised mice. **(F)** Plasma concentrations of PAA, PAGln, and PAGly in WT- and ΔBT0430-colonised mice. **(G)** Body weight, liver/body weight ratio, and fat/body weight ratio. **(H)** Non-fasted blood glucose concentrations. **(I)** Plasma ALT and AST activities. **(J)** Composite SAF score, steatosis score, activity score, and fibrosis score. **(K)** Representative histological images of liver sections stained with haematoxylin and eosin (H&E) and picrosirius red (PSR). Scale bars, 50 µm. **(L)** Principal component analysis of liver RNA-seq data from WT- and ΔBT0430-colonised mice using variance-stabilised counts. Each point represents one animal; dashed ellipses indicate 95% confidence regions. **(M)** Volcano plot of differential gene expression in ΔBT0430-versus WT-colonised livers. Red and blue points indicate significantly upregulated and downregulated genes, respectively (adjusted *P* < 0.05); selected genes are labelled. **(N)** Overlap between genes upregulated by PAGln supplementation and genes downregulated in ΔBT0430-colonised livers (adjusted *P* < 0.05). Significance was assessed using a hypergeometric test. **(O)** Log2 fold changes of the eight shared genes in the PAGln versus vehicle and ΔBT0430 versus WT contrasts. Data are represented as mean ± SEM. For (E, G-J), n = 10 (WT) and 8 (ΔBT0430); for (F), n = 5 per group; and for (L-O), n = 6 per group. Statistical significance was determined by two-way ANOVA for (C); unpaired two-tailed t-test for (E-I); Mann-Whitney test for (J); and for (L-O), by Wald test with Benjamini-Hochberg correction in DESeq2, with overlap significance in (N) assessed by hypergeometric test. *p < 0.05, **p < 0.01, ***p < 0.001.

To determine the contribution of BT0430 to MASLD in vivo, germ-free mice were mono-colonised with wild-type or ΔBT0430 *B. thetaiotaomicron* and maintained on the LID21 diet for 12 weeks (**Figure 5D**). Faecal bacterial 16S rRNA gene copy numbers did not differ significantly between groups, indicating comparable bacterial colonisation, whereas BT0430 DNA was undetectable in ΔBT0430-colonised mice, confirming maintenance of the deletion in vivo (**Figure 5E**). Plasma PAGly concentrations were markedly reduced in ΔBT0430-colonised mice, with a similar trend for PAGln, whereas PAA concentrations were unchanged (**Figure 5F**).

We next examined the metabolic and hepatic consequences of BT0430 deletion. ΔBT0430 colonisation did not significantly alter body weight and showed a trend toward a lower liver-to-body weight ratio, but significantly reduced the fat-to-body weight ratio compared with wild-type colonisation (**Figure 5G**). Non-fasted blood glucose was also significantly lower in ΔBT0430-colonised mice (**Figure 5H**). Plasma ALT activity showed a trend toward reduction, whereas AST activity did not differ significantly between groups (**Figure 5I**). Histological assessment showed significantly lower composite SAF, steatosis, and fibrosis scores in ΔBT0430-colonised mice, whereas the activity score did not differ significantly (**Figure 5J-K**).

We next performed liver RNA sequencing to characterise the hepatic transcriptional changes associated with BT0430 deletion. Principal component analysis showed partial separation between wild-type and ΔBT0430-colonised mice along PC1, which accounted for 64% of the variance (**Figure 5L**). Differential expression analysis identified 119 upregulated and 77 downregulated genes in ΔBT0430-colonised livers (**Figure 5M**). Intersecting the 183 genes induced by PAGln supplementation with the 77 genes reduced in ΔBT0430-colonised livers identified eight shared genes, an overlap greater than expected by chance (hypergeometric P = 1.6 × 10⁻⁵; **Figure 5N**). All eight genes changed in opposite directions across the two experiments, increasing with PAGln supplementation and decreasing following BT0430 deletion (**Figure 5O**). These included the detoxification and transport genes *Por* and *Abcc4*, the stress hormone and insulin signalling genes *Fkbp5* and *Zbtb16*, the proliferation-associated genes *Smc4* and *Tfap4*, and *Tbc1d8* and *Irf2bpl*.

Together, these findings establish BT0430 as a bacterial determinant of PAA production by *B. thetaiotaomicron* and show that its deletion reduces circulating PAA conjugates and ameliorates hepatic steatosis and fibrosis in gnotobiotic mice.

### Microbiota-derived phenylalanine metabolites are elevated in the cardiometabolic subtype of human MASLD

Having shown that microbial phenylalanine metabolism contributes to MASLD in mice, we next asked whether related circulating metabolites distinguish clinically relevant subtypes of human MASLD. We used a previously validated, data-driven clustering framework in which 1,389 participants with obesity from the ABOS cohort were partitioned into six clusters by partitioning-around-medoids clustering based on six routine clinical variables: age, BMI, LDL cholesterol, ALT, HbA1c, and triglycerides^28^. Liver biopsy identified two clusters enriched for MASH, which were labelled according to their dominant clinical features: a cardiometabolic MASLD (CM) cluster, characterised by an increased prevalence of type 2 diabetes, dyslipidaemia and hypertension, and a liver-specific MASLD (LS) cluster, characterised by elevated ALT and AST. The remaining four clusters were pooled into a non-MASH reference group (**Figure 6A**). Using plasma metabolomics data from this cohort together with its published cluster assignments, we examined plasma levels of PAA, PAGln and PCS across the control, CM and LS groups.

**Figure 6.**
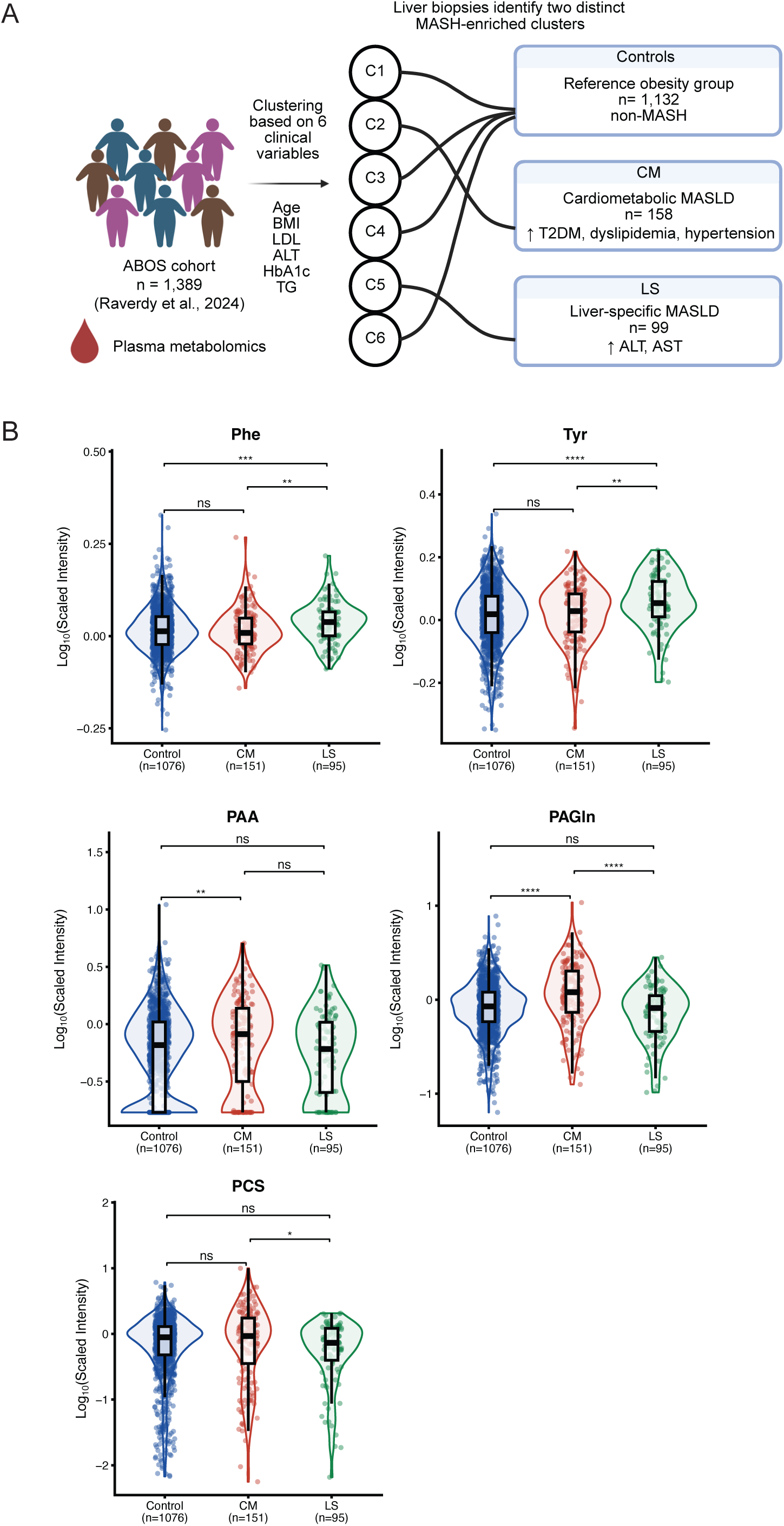
Circulating PAGln is elevated in cardiometabolic MASLD in humans. **(A)** Schematic of the previously established MASLD subtypes in the ABOS cohort (n = 1,389 individuals with obesity; Raverdy *et al*.). Participants were partitioned into six clusters (C1 to C6) by partitioning-around-medoids clustering based on six clinical variables (age, BMI, LDL cholesterol, ALT, HbA1c, and triglycerides). Two clusters enriched for MASH were classified as cardiometabolic MASLD (CM; n = 158), characterised by a higher prevalence of type 2 diabetes, dyslipidaemia, and hypertension, or liver-specific MASLD (LS; n = 99), characterised by elevated ALT and AST. The remaining clusters comprised the reference group (Controls, n = 1,132). **(B)** Plasma metabolite intensities of phenylacetic acid (PAA), phenylacetylglutamine (PAGln), p-cresol sulfate (PCS), phenylalanine (Phe), and tyrosine (Tyr) from reanalysis of ABOS plasma metabolomics data, stratified according to the previously established MASLD subtypes (Controls, n = 1,076; CM, n = 151; LS, n = 95). Values are shown as log10-transformed, scaled intensities. Each point represents an individual participant; violin plots show the distribution with overlaid boxplots. Statistical significance was determined using the Kruskal–Wallis test with Dunn’s post hoc test. *P < 0.05, **P < 0.01, ****P < 0.0001; ns, not significant.

PAA was significantly higher in the CM cluster than in controls, whereas levels in the LS cluster did not differ significantly from either group. PAGln was markedly higher in the CM cluster than in both controls and the LS cluster, while LS and control levels were comparable. PCS did not differ significantly between the CM cluster and controls or between the LS cluster and controls, although levels were modestly higher in the CM than in the LS cluster (**Figure 6B**). In contrast, plasma phenylalanine and tyrosine were elevated in the LS cluster compared with the other groups (**Figure 6B**).

Together, these findings indicate that circulating PAGln is selectively elevated in the cardiometabolic subtype of human MASLD, suggesting a potential link between this microbial metabolic pathway and the cardiometabolic phenotype of MASLD.

## Discussion

This study demonstrates that the contribution of the gut microbiota to MASLD is strongly influenced by dietary protein availability. Through caecal and plasma metabolomics, we identified microbial phenylalanine metabolism as a candidate pathway underlying this diet-dependent effect. PAGln supplementation exacerbated hepatic steatosis and liver injury without altering adiposity or systemic glucose homeostasis, whereas deletion of a phenylpyruvate ferredoxin oxidoreductase in *B. thetaiotaomicron* disrupted PAA production in vitro, reduced circulating PAGln/PAGly, and ameliorated hepatic steatosis and fibrosis in mono-colonised germ-free mice. Together, these findings support a model in which dietary protein availability modulates the contribution of microbial phenylalanine metabolism to MASLD.

Our findings connect previously established relationships between dietary protein, microbial amino acid metabolism, and MASLD. Higher animal protein intake has been associated with increased odds of MASLD and histological disease activity^11,12^, while dietary amino acids can contribute directly to hepatic lipogenesis.^13^ In parallel, dietary protein reaching the colon undergoes microbial fermentation, generating aromatic amino acid catabolites and other metabolites.^29–32^ Gut microbial communities can also transmit susceptibility to hepatic steatosis.^33,20,22^ Our data suggest that these relationships are interconnected: microbiota depletion improved hepatic outcomes under LID21 but not LID7 conditions. This does not imply that the microbiota is required for MASLD, as disease remained evident under LID7, but rather that dietary context may modify the contribution of specific microbial metabolic functions to disease. Such diet dependence may also contribute to heterogeneity in microbiome associations reported across MASLD cohorts.^34,35^

Our genetic experiments provide functional evidence linking a defined bacterial metabolic activity to hepatic disease. Previous studies have used bacterial genetics to assign circulating aromatic amino acid metabolites to specific microbial pathways,^36,37^ and more recently identified distinct routes to PAA production, including *porA*-dependent metabolism in *Clostridium sporogenes* and phenylpyruvate ferredoxin oxidoreductase loci in *B. thetaiotaomicron*.^19^ Here, deletion of BT0430, encoding a phenylpyruvate ferredoxin oxidoreductase in *B. thetaiotaomicron*, abolished detectable PAA production under the conditions tested in vitro and reduced circulating PAGln/PAGly in vivo, while ameliorating hepatic steatosis and fibrosis. These findings extend previous work linking bacterial genes to metabolite production by demonstrating that perturbation of a defined microbial PAA-producing function can alter a histologically defined disease phenotype. The predominance of PAGly rather than PAGln in mice is consistent with species-specific host conjugation of PAA, whereby mice favour glycine conjugation whereas glutamine conjugation predominates in humans.^19^ Thus, microbial PAA production represents a metabolic step upstream of species-dependent host conjugation. However, the contribution of this individual pathway within a complex microbial community, in which multiple taxa and enzymatic routes can contribute to PAA production, remains to be determined.

The liver transcriptomic analyses provided insight into the host response associated with this pathway. PAGln supplementation was accompanied by enrichment of interferon α and γ responses, IL-2/STAT5 signalling, complement, TGF-β signalling and E2F targets, whereas no lipid metabolism-related Hallmark gene set was significantly enriched. Induction of the STAT target genes *Cish* and *Socs3*, together with reduced *Bcl6*, was consistent with altered STAT-associated signalling. PAGln failed to directly increase triglyceride accumulation in cultured hepatocytes, suggesting that its in vivo effects may involve pathways not captured by this simplified hepatocyte model. Comparison of the two liver transcriptomic datasets identified eight genes induced by PAGln and reduced following disruption of bacterial PAA production, an overlap greater than expected by chance. Among these, *Fkbp5* is of particular interest because FKBP51 can modulate insulin signalling through PHLPP-mediated regulation of AKT2, and *Fkbp5*-deficient mice show resistance to diet-induced metabolic dysfunction.^38^ Nevertheless, the overlap represents only a small subset of PAGln-responsive genes, and these transcriptional findings should therefore be considered hypothesis-generating. PAGln has previously been shown to signal through adrenergic receptors in cardiovascular contexts,^17^ but its molecular targets and relevant target tissues in MASLD remain to be determined.

The human data further suggest that the relevance of microbial phenylalanine metabolism may differ across MASLD phenotypes. Circulating PAGln was selectively elevated in the cardiometabolic MASLD subtype, whereas phenylalanine and tyrosine were higher in the liver-specific subtype. This distinction is notable given previous associations of PAGln with cardiovascular and metabolic disease.^17,18,19^ However, the human analysis is cross-sectional and does not establish whether increased PAGln contributes to cardiometabolic MASLD or instead reflects metabolic, dietary, renal, or microbial features associated with this phenotype. Prospective studies incorporating dietary information and microbiome profiling will be required to determine whether microbial PAA production identifies a biologically distinct subset of MASLD.

Future studies should define the molecular targets and target tissues through which PAGln and PAGly influence host metabolism, establish the contributions of distinct microbial PAA-producing pathways within complex microbial communities, and determine whether dietary protein intake stratifies the relevance of this pathway in human MASLD. Such studies may clarify whether diet- or microbiota-directed interventions targeting microbial phenylalanine metabolism could benefit specific metabolic subtypes of MASLD.

## Methods

### Mice and animal experiments

Male C57BL/6J and C57BL/6N mice were obtained from either the Animal Research Facility (ARF) at the Lee Kong Chian School of Medicine, Nanyang Technological University, Singapore, or InVivos Pte Ltd, and were maintained under specific-pathogen-free (SPF) conditions at the ARF. Germ-free C57BL/6N mice were obtained and maintained in laminar-flow isolators in the germ-free facility at the ARF. Mice were housed under a 12-h light-dark cycle (lights on from 7 am to 7 pm) with ad libitum access to food and water. For experiments using the Liver Disease Progression Aggravation Diet (LIDPAD) model^25^, mice were housed under thermoneutral conditions (30°C).

The MASLD-inducing diets were adapted from the LIDPAD model with modifications to the protein content. The percentages of kcal derived from protein were 21% for LID21, 11% for LIDPAD, and 7% for LID7. Unless otherwise specified, dietary interventions were initiated at 8 weeks of age and continued for 12 weeks. The nutritional composition of all diets is listed in **Supplementary Table 2**.

For antibiotic treatment, mice were given ampicillin (Sigma, A9393; 1 g/L), vancomycin (Axil Scientific, V-200-5; 0.5 g/L), neomycin (Sigma, N6386; 1 g/L), or metronidazole (Sigma, M3761; 1 g/L) in their drinking water. When indicated, a combination of these antibiotics was administered in their drinking water to broadly deplete the gut microbiota.

For metabolite supplementation experiments, 8-week-old male C57BL/6J mice were maintained on the LIDPAD diet under thermoneutral conditions (30°C) for 12 weeks. For PAGln supplementation, mice were randomly assigned to vehicle or phenylacetylglutamine (PAGln; TCI Chemicals, P2989; 100 mg/kg) groups (n = 10 per group) and administered the respective treatment daily by intraperitoneal injection. One mouse from the PAGln group died before the end of the experiment. In a separate cohort, mice were randomly assigned to control or p-cresol sulfate (PCS; TCI Chemicals, P2091) groups (n = 10 per group). PCS was administered at 0.05% (w/v) in the drinking water throughout the 12-week intervention, whereas control mice received unsupplemented drinking water. Drinking water was freshly prepared and replaced every 2 days, and water intake was recorded throughout the intervention. For gnotobiotic experiments, GF C57BL/6N mice were orally administered 200 μL of either wild-type or ΔBT0430 *B. thetaiotaomicron*. Successful colonisation was confirmed by quantitative PCR of bacterial 16S rRNA genes in faecal samples.

All animal procedures were approved by, and performed in accordance with, the Nanyang Technological University Institutional Animal Care and Use Committee (NTU-IACUC A22094).

### Biochemistry analysis

Plasma AST and ALT activities were measured using the IDEXX Catalyst One Veterinary Chemistry Analyser.

### Glucose and insulin tolerance tests

For glucose and insulin tolerance tests, mice were fasted for 5 h. For oral glucose tolerance tests (OGTT), glucose was administered at 2 g kg⁻¹ body weight by oral gavage. For insulin tolerance tests (ITT), insulin was administered at 0.75 U kg⁻¹ body weight by intraperitoneal injection. Blood glucose was measured using a glucometer (Accu-Chek Performa) in blood samples collected from the tail at −30, 0, 15, 30, 60 and 90 min for ITT, and at −30, 0, 15, 30, 60, 90 and 120 min for OGTT.

### Flow cytometry

Peripheral blood was collected from the retro-orbital plexus using heparin-coated capillary tubes. Erythrocytes were lysed in 1× BD Pharm Lyse™ lysing buffer on ice for 5 min and washed twice before staining. Cells obtained from 100 µL of lysed blood were incubated with Fc Block (BD Biosciences) for 10 min, followed by staining with the antibodies listed in **Supplementary Table 4** for 30 min at 4 °C. Cells were subsequently fixed with BD Cytofix before acquisition. Data were acquired on an LSRFortessa X-20 flow cytometer (BD Biosciences) and analysed using FlowJo v10. Gating strategies for whole blood are shown in **Supplementary Figure 3A**.

### Histopathology and liver scoring

Liver samples were fixed in 10% neutral-buffered formalin for 24 h, paraffin-embedded, sectioned at 5 μm, and stained with hematoxylin and eosin or Picrosirius Red. Three blinded assessors independently evaluated steatosis, activity, and fibrosis using the SAF scoring system, and the scores were averaged for subsequent analyses. Steatosis was graded on a scale of 0-3 according to the proportion of hepatocytes containing lipid droplets: S0 for under 5%, S1 for 5-33%, S2 for 33-66%, and S3 for over 66%. The activity score (0-4) was calculated as the sum of hepatocyte ballooning (0-2) and lobular inflammation (0-2). Hepatocyte ballooning was scored as 0 for normal hepatocytes, 1 for clusters of hepatocytes with a rounded shape and pale cytoplasm, and 2 for enlarged hepatocytes with clear cytoplasm. Lobular inflammation was scored as 0 for no inflammatory foci, 1 for ≤2 foci per ×20 field, and 2 for >2 foci per ×20 field. Fibrosis was staged from F0 to F4: F0, no fibrosis; F1, perisinusoidal fibrosis in zone 3 (F1a or F1b) or periportal fibrosis (F1c); F2, perisinusoidal and periportal fibrosis; F3, bridging fibrosis; and F4, cirrhosis.

### Cell culture

AML12 cells were maintained in DMEM supplemented with 10% (v/v) foetal bovine serum and 1% (v/v) penicillin-streptomycin at 37 °C in a humidified atmosphere of 5% CO₂, and were passaged every 3 days. For treatment experiments, cells were seeded at 1 × 10⁵ cells per well in 24-well plates or 5 × 10⁵ cells per well in 6-well plates and cultured for 24 h prior to treatment. The medium was then replaced with fresh DMEM containing the indicated treatments: DMEM alone (control), 10-1000 µM *p*-cresol sulfate (P2091, TCI Chemicals), phenylacetylglycine (P0131, TCI Chemicals) or phenylacetylglutamine (P2989, TCI Chemicals). Cells were treated for 24 h before downstream analysis.

To model lipid overload, we performed a parallel set of experiments using an oleic acid–palmitic acid mixture (OAPA). Oleic acid (Sigma, O1383) and palmitic acid (Sigma, P0500) were combined at a 2:1 molar ratio (133.3 µM oleic acid and 66.7 µM palmitic acid; 200 µM total) and conjugated to fatty-acid-free BSA. Cells were co-treated with OAPA and 100 µM p-cresol sulfate, phenylacetylglycine, or phenylacetylglutamine for 24 h, with BSA vehicle serving as the lipid-free control, prior to downstream analysis.

### Triglyceride quantification

Intracellular triglyceride levels were measured using the Triglyceride Quantification Assay Kit (Abcam, ab65336). Briefly, treated cells were washed with cold PBS and lysed in 5% Nonidet P-40, followed by heat treatment to fully solubilise triglycerides. Cleared lysates were diluted tenfold and assayed in duplicate in a fluorometric format according to the manufacturer’s instructions. After incubation with cholesterol esterase and reaction mix, fluorescence was measured at Ex/Em 535/587 nm. Total protein concentration from the same lysates was quantified using the Pierce BCA Protein Assay Kit (Thermo Fisher Scientific, 10678484), and triglyceride levels were normalised to protein content.

### Bacterial strains

*Bacteroides thetaiotaomicron* VPI-5482 and the *B. thetaiotaomicron* Δ*BT0430* mutant strain generated from the parental VPI-5482 strain were used in this study. All bacterial strains were cultured anaerobically at 37°C in BHIS medium, consisting of brain heart infusion medium supplemented with vitamin K1 (1 mg/L), hemin (1.2 mg/L), and L-cysteine (0.5 g/L). Anaerobic cultivation was performed in an anaerobic chamber with a gas mixture of 5% hydrogen, 10% carbon dioxide, and 85% nitrogen. Bacterial cultures were grown under static conditions without shaking.

### *B. thetaiotaomicron* ΔBT0430 mutant generation

A BT0430 deletion mutant was generated in *B. thetaiotaomicron* VPI-5482 by homologous recombination using the shuttle plasmid pLGB13 carrying homologous arms flanking the target locus^39^. The plasmid was introduced into E. coli S17 λpir and mobilised into *B. thetaiotaomicron* by conjugation. Recombinants were selected using erythromycin (Sigma, E5389) and gentamicin (Gibco, 15710064), followed by counterselection on agar supplemented with anhydrotetracycline (ATC) (Sigma, 37919) to isolate double-crossover mutants. Candidate ΔBT0430 clones were identified by growth on ATC plates and loss of growth on erythromycin and gentamicin plates. Successful deletion of BT0430 was verified by PCR and whole-genome sequencing. All primers, including the primers used to amplify homologous arms and to check identities of mutants, are included in **Supplementary Table 3**.

### Quantification of phenylacetic acid using LC-MS

Proteins were precipitated by treating 50 µL of sample with 200 µL of ice-cold methanol. Following centrifugation at 14,000 x g for 15 min at 4 °C, 50 µL of supernatant or external standard was derivatised by adding 25 µL of 250 mM 3-nitrophenylhydrazine (3-NPH) in 75% methanol and 25 µL of 150 mM 1-ethyl-3-(3-dimethylaminopropyl)carbodiimide (EDC) in 75% methanol containing 7.5% pyridine, with incubation for 30 min at 40 °C^40^. Quantitative analysis was performed on a Waters UPLC system coupled to a triple quadrupole mass spectrometer with an electrospray ionisation source (Waters, MA, USA), with the derivatised analyte separated on an Acquity UPLC BEH C18 column (2.1 mm × 100 mm, 1.7 µm; Waters) using mobile phases of water (A) and acetonitrile (B), each containing 0.01% formic acid, delivered at 0.6 mL/min with the following gradient: 14% B (0–1 min), 14–20% B (1–2 min), 20–28% B (2–5 min), 28–40% B (5–7 min), 40–100% B (7–8 min) and 100% B (8–10 min); the column temperature was 40 °C and the injection volume 5 µL. Detection was carried out in negative mode by multiple reaction monitoring (capillary voltage, 2,500 V; source temperature, 150 °C; desolvation temperature, 500 °C), monitoring phenylacetic acid at the transition m/z 236.0 → 137.0 (fragmentor voltage, 20 V; collision energy, 18 eV). Peak areas were integrated using TargetLynx XS software (Waters), and concentrations were quantified against calibration curves prepared from external standards.

### Targeted metabolomics analysis

Targeted metabolomic profiling of mouse plasma was performed using a 300-multiple reaction monitoring panel. Plasma samples (80 μL) were extracted with 320 μL ice-cold methanol containing internal standards, vortexed, sonicated in an ice-water bath for 15 min, incubated at −40 °C for 1 h, and centrifuged at 12,000 rpm for 15 min at 4 °C. Supernatants were collected, and 1 μL was subjected to UHPLC–MS/MS analysis using an Agilent 1290 Infinity II system coupled to a SCIEX 6500 QTRAP mass spectrometer.

Metabolites were separated on an ACQUITY UPLC BEH C18 column (2.1 × 150 mm, 1.7 μm) using 0.1% formic acid in water as mobile phase A and 95:5 methanol:water containing 10 mM ammonium formate as mobile phase B. The autosampler was maintained at 6 °C. Mass spectrometric analysis was performed using an IonDrive Turbo V electrospray ionization source with the following settings: curtain gas, 35 psi; ion spray voltage, 5,000 V in positive mode and −4,500 V in negative mode; source temperature, 450 °C; and GS1 and GS2, 50 psi. Data acquisition and quantification were performed using SCIEX Analyst v1.7.3 and BIOTREE Bio-Bud v2.0.3.

Metabolite concentrations were determined using calibration curves with R^2^ ≥ 0.95. Pooled quality control samples were injected after every 10 samples to monitor analytical stability. Internal standard relative standard deviation was <20% and retention time variation was <2 s. Final concentrations were corrected for extraction volume and reported in nmol/L.

### Untargeted metabolomics analysis of caecal content

Frozen mouse caecal contents (∼25 mg) were extracted in 500 μL methanol:acetonitrile:water (2:2:1, v/v/v) containing deuterated internal standards for analysis of polar and non-polar metabolites. Samples were vortexed, homogenized at 35 Hz for 4 min for three cycles, sonicated for 5 min in a 4 °C water bath for three cycles, incubated at −40 °C for 1 h, and centrifuged at 13,800 x g for 15 min at 4 °C. Supernatants were collected for LC–MS/MS analysis. Pooled quality control samples were prepared by combining aliquots from all samples.

Polar metabolites were separated on an ACQUITY UPLC BEH Amide column (2.1 × 100 mm, 1.7 μm, Waters) using a Vanquish UHPLC system (Thermo Fisher Scientific), with 25 mM ammonium acetate and 0.25% ammonium hydroxide in water as mobile phase A and acetonitrile as mobile phase B. Non-polar metabolites were separated on a Kinetex C18 column (2.1 × 100 mm, 2.6 μm, Phenomenex), with 0.01% acetic acid in water as mobile phase A and isopropanol:acetonitrile (1:1, v/v) as mobile phase B. Mass spectrometric detection was performed on an Orbitrap Exploris 120 mass spectrometer (Thermo Fisher Scientific) operated in positive and negative electrospray ionization modes, with resolutions of 60,000 for MS1 scans and 15,000 for MS/MS scans using information-dependent acquisition.

Raw data were converted to mzXML format and processed using an in-house R-based pipeline built on XCMS for peak detection, retention time alignment and peak integration. Metabolites were annotated using BiotreeDB v3.07 on the basis of MS1, MS2 and retention time matching according to Metabolomics Standards Initiative criteria. Data were filtered on the basis of relative standard deviation, missing values were imputed with half of the minimum detected value, and signal intensities were normalized to internal standards.

### K-means clustering analysis

Untargeted metabolomics data from caecal contents were analysed to identify metabolic pathways associated with the protein- and microbiota-dependent phenotype. Annotated metabolites were subjected to k-means clustering based on their intensity profiles across the four experimental groups using the kmeans function in base R (stats package; Hartigan–Wong algorithm; iter.max = 200, nstart = 25). For exploratory pattern discovery, metabolites were partitioned into nine clusters using a prespecified k = 9. Clusters 4 and 7, comprising 1,066 and 765 metabolites, respectively, were selected for downstream analysis because their abundance profiles most closely matched the protein- and microbiota-dependent pattern observed in the phenotypic data, with higher abundance under protein-sufficient, microbiota-intact conditions and lower abundance following microbiota depletion and/or protein restriction. Within each cluster, metabolites significantly altered across experimental conditions were identified using linear modelling with the limma package. Significant metabolites from clusters 4 and 7 were then analysed separately using the pathway analysis module in MetaboAnalyst 6.0. Pathway enrichment and pathway impact were calculated and visualised as plots of pathway significance versus pathway impact.

### Differential abundance and pathway analysis

Differential abundance was assessed with limma using a two-factor design (∼ Protein × ABX), with contrasts for the average microbiota effect, the protein effect within each microbiota state, and their interaction; variance was moderated by empirical Bayes and P values adjusted by the Benjamini-Hochberg method. Metabolites were classified as showing a microbiota-dependent protein response if significant for all three contrasts (FDR < 0.05) and if the magnitude of the protein effect was reduced after microbiota depletion. Pathway testing was performed without prior selection of pathways: KEGG membership was retrieved using KEGGREST for the 1,171 metabolites carrying KEGG identifiers, and all 138 pathways containing 5 to 500 metabolites were tested using both a competitive test (camera) and a self-contained rotation test (mroast, 9,999 rotations), with FDR applied across pathways within each contrast. Metabolites were additionally ranked by moderated t-statistic across the full dataset; reported percentiles denote position within this ranking. Analyses were repeated using level 1 annotations only as a sensitivity analysis.

### Faecal DNA extraction and quantification

Faecal DNA was extracted using the bead-beating method as previously described^41^. Briefly, a frozen faecal pellet was suspended in 500 µL of extraction buffer (200 mM Tris (pH 8.0), 200 mM NaCl, 20 mM EDTA), 210 µL of 20% SDS, 500 μL of a mixture of phenol:chloroform:isoamyl alcohol (25:24:1, pH 7.9), and 500 μL of 0.1 mm diameter zirconia/silica beads (BioSpec Products). Microbial cells were lysed by mechanical disruption with a homogenizer (TissueLyser II, Qiagen) for 5 minutes at room temperature, followed by centrifugation to collect the supernatant and precipitation with isopropanol. DNA was then purified with a PCR purification kit (Qiagen) and quantified using NanoDrop and Qubit analysers.

### Faecal metagenomics

Faecal DNA from mice treated with individual antibiotics was submitted to GENEWIZ for shotgun metagenomic sequencing. After DNA quality control using Qubit and gel electrophoresis, libraries were constructed by fragmentation, end repair, adaptor ligation, size selection, and index PCR, and sequenced on an Illumina NovaSeq platform in paired-end 2 × 150 bp mode. Raw sequencing data were processed using bcl2fastq (v2.17.1.14) to generate pass-filter reads, which were further quality filtered using cutadapt (v1.9.1) by removing adaptor sequences and low-quality reads, excluding reads with an N base ratio greater than 10%, trimming bases with terminal quality scores <20, and retaining reads with a minimum length of 75 bp. Clean reads were assembled using MEGAHIT (v1.2.9), coding sequences were predicted using Prodigal (v3.02), and redundant genes were clustered using MMseq2 at 95% sequence identity and 95% coverage to generate a non-redundant unigene catalogue. Reads were then mapped back to the unigene catalogue using SOAPAligner (v2.21) to estimate gene abundance normalised to gene length. Taxonomic annotation was assigned based on the NR database using BLAST (version 2.2.31+), and the resulting taxa abundance and gene abundance tables were used for downstream analyses described in this paper. KEGG orthologs were annotated based on the KEGG database.^42^

All downstream bioinformatic analyses were performed in R (version 4.6.0) using taxonomic and gene abundance tables generated from shotgun metagenomic sequencing. Alpha diversity was assessed using the Chao1, Shannon and Simpson indices with the phyloseq and vegan packages. Beta diversity was evaluated by principal coordinates analysis (PCoA) based on Bray–Curtis distances using vegan, and taxonomic composition was summarised as relative abundance at the relevant taxonomic levels. Associations between bacterial taxa and MASLD severity were assessed using MaAsLin2. Taxa associated with plasma ALT activity or SAF score were identified at a Benjamini–Hochberg-adjusted q < 0.05, and taxa associated with both parameters were selected for further analysis. To examine microbial functions potentially involved in PAA production, candidate 2-oxoacid:ferredoxin oxidoreductase gene families (*porA–D*, *korA–D*, *iorA/B*, *vorD*, and *por/nifJ*) were identified from the metagenomic gene abundance data. Their abundances were correlated with plasma ALT and AST activities and histological SAF, steatosis, activity and fibrosis scores using Spearman’s rank correlation. The presence of these candidate genes was also examined across the taxa associated with both ALT and SAF score. Results were visualised using ggplot2 and pheatmap.

### Bacterial absolute quantification

Total bacterial load was quantified by qPCR targeting the 16S rRNA gene, using genomic DNA extracted from faecal or caecal samples as described above. A standard curve was generated from serial ten-fold dilutions of bacterial genomic DNA at known copy concentrations, allowing absolute quantification of 16S rRNA gene copies per reaction. Bacterial load is expressed as 16S rRNA gene copies per mg of faeces, using universal 16S rRNA primers [331F: TCCTACGGGAGGCAGCAGT, 797R: GGACTACCAGGGTATCTAATCCTGTT], with thermal cycling conditions of initial denaturation (95 °C, 30 s), followed by 40 cycles of denaturation (95 °C, 15 s) and annealing/extension (60 °C, 30 s), and melt-curve analysis (65–95 °C, 0.5 °C increments) to confirm amplicon specificity. No-template controls were included in every run.

### RNA extraction, library preparation and sequencing

Total RNA was extracted from snap-frozen liver tissue using the RNeasy Mini Kit (Qiagen) according to the manufacturer’s instructions. RNA integrity and concentration were assessed before library preparation. Library construction and sequencing were performed by BGI Genomics. Poly(A)-containing mRNA was enriched from total RNA using oligo(dT)-conjugated magnetic beads and fragmented, and first-strand cDNA was synthesised using random N6 primers. Second-strand synthesis incorporated dUTP to preserve strand information. Double-stranded cDNA was end-repaired, 5’-phosphorylated and A-tailed, followed by ligation of bubble-shaped adapters. Ligation products were amplified by PCR, during which the dUTP-marked strand was degraded by uracil-DNA glycosylase, and the products were denatured and circularised to generate single-stranded circular DNA libraries. Libraries were converted to DNA nanoballs and sequenced on the DNBSEQ-G400 platform with 150 bp paired-end reads.

### Read processing, alignment and quantification

Raw reads were filtered with SOAPnuke (v1.5.6) to remove adapter-containing reads, reads with more than 0.1% unknown bases, and low-quality reads, defined as those in which more than 20% of bases had a quality score below 15. Clean reads were aligned to the *Mus musculus* GRCm39 reference genome (NCBI GCF_000001635.27, annotation v2201) using HISAT2 (v2.2.1) with strand-specific settings appropriate to the dUTP protocol. Mean total genome mapping rate was 98.54%. For quantification, clean reads were aligned to the reference transcriptome using Bowtie2 (v2.5.0) and expected counts and TPM values were estimated with RSEM (v1.3.1). A total of 17,319 genes were detected across the dataset.

### Differential expression analysis

RSEM expected counts were rounded to integers and imported into R for differential expression analysis using DESeq2 (v1.40.2). The two contrasts, PAGln versus vehicle and ΔBT0430 versus wild-type colonisation, were analysed independently. Size factors were estimated using the median-of-ratios method, and differential expression was assessed using the Wald test with DESeq2 independent filtering. *P* values were adjusted using the Benjamini–Hochberg procedure, and genes with adjusted *P* < 0.05 were considered differentially expressed. For volcano plots, log2 fold changes were shrunk using the apeglm estimator, while unshrunken adjusted *P* values were used to determine statistical significance. For principal component analysis and heatmaps, counts were variance-stabilising transformed, and heatmap values were displayed as per-gene z-scores of transformed expression.

### Gene set enrichment analysis

Gene set enrichment analysis (GSEA) was performed using fgsea (v1.38.0) against the mouse Hallmark gene set collection from the Molecular Signatures Database (MSigDB). All expressed genes were ranked by signed significance, calculated as sign(log2 fold change) × −log10(*P*) from the DESeq2 results. Enrichment significance was assessed using 10,000 permutations, with *P* values adjusted using the Benjamini–Hochberg procedure.

### Sensitivity analysis

Sample-level diagnostics identified one PAGln-treated sample (N3) as a candidate outlier based on principal component analysis, pairwise Pearson correlations, and expression density distributions. The sample was retained in the primary analysis, and the complete differential expression and GSEA pipeline was repeated after excluding N3 as a sensitivity analysis. Hallmark gene sets were considered robust when they remained significantly enriched in the same direction in both analyses. Cook’s distances were additionally examined to assess sample-level influence on differential expression results.

### Cross-contrast overlap

Overlap between genes induced by PAGln supplementation and those reduced in ΔBT0430-colonised livers was assessed using a hypergeometric test. The background universe comprised genes tested in both differential expression contrasts rather than the whole annotated genome.

## Statistical analyses

Statistical analyses were performed using R (v4.6.0) and GraphPad Prism 9. Data are presented as mean ± s.e.m. Comparisons between two groups were performed using unpaired two-tailed Student’s t-tests or Mann–Whitney tests, as appropriate. Comparisons among multiple groups were performed using one- or two-way ANOVA with appropriate post hoc multiple-comparison tests or Kruskal–Wallis tests with Dunn’s multiple-comparison test, as indicated in the corresponding figure legends. Associations between plasma metabolite abundance and MASLD severity parameters, including ALT, AST, SAF score, steatosis grade, activity score and fibrosis stage, were assessed using Spearman’s rank correlation. Differential metabolite analysis was performed using the limma package, and differential gene expression analysis was performed using DESeq2 as described above. A two-sided P < 0.05 was considered statistically significant unless otherwise specified. For analyses involving multiple testing, significance was assessed using Benjamini–Hochberg-adjusted *P* values or *q* values, as indicated.

## Data availability

The metagenomic and RNA-seq datasets generated in this study have been deposited in the NCBI Sequence Read Archive (SRA) under BioProject accession numbers PRJNA1521042 and PRJNA1528863, respectively. The human plasma metabolomics data analysed in this study are publicly available through BioStudies under accession S-BSST1479 from the previously published study by Raverdy *et al*.^28^

## Acknowledgements

This research is supported by the following: Vascular Research Initiative, Strategic Academic Initiative 022415-00001 (to K.K. and N.S.T.); Singapore Ministry of Education under its Singapore Ministry of Education Academic Research Fund Tier 2 (MOE-T2EP30223-0008) and Tier 1(RG125/25); Singapore Ministry of Health’s National Medical Research Council (CS-IRG MOH-001997). We gratefully acknowledge A/Prof Christine Cheung at Nanyang Technological University for her leadership of the Vascular Research Initiative and her support of this research.

## Author contributions

Y.R.B. and K.K. conceived and designed the study. Y.R.B. performed the majority of the experiments, analysed and interpreted the data, and prepared the original draft of the manuscript. D.N.A.B.M. and F.E.S. assisted with experiments and data collection. T.E., S.H.W., R.D., W.-K.W., N.S.T. and T.W. contributed to the study through scientific collaboration, discussion, and interpretation of the results. K.K. supervised the study, provided overall scientific direction and guidance, interpreted the data, and revised the manuscript. All authors reviewed and edited the manuscript and approved the final version.

---

**Supplementary Figure 1.**
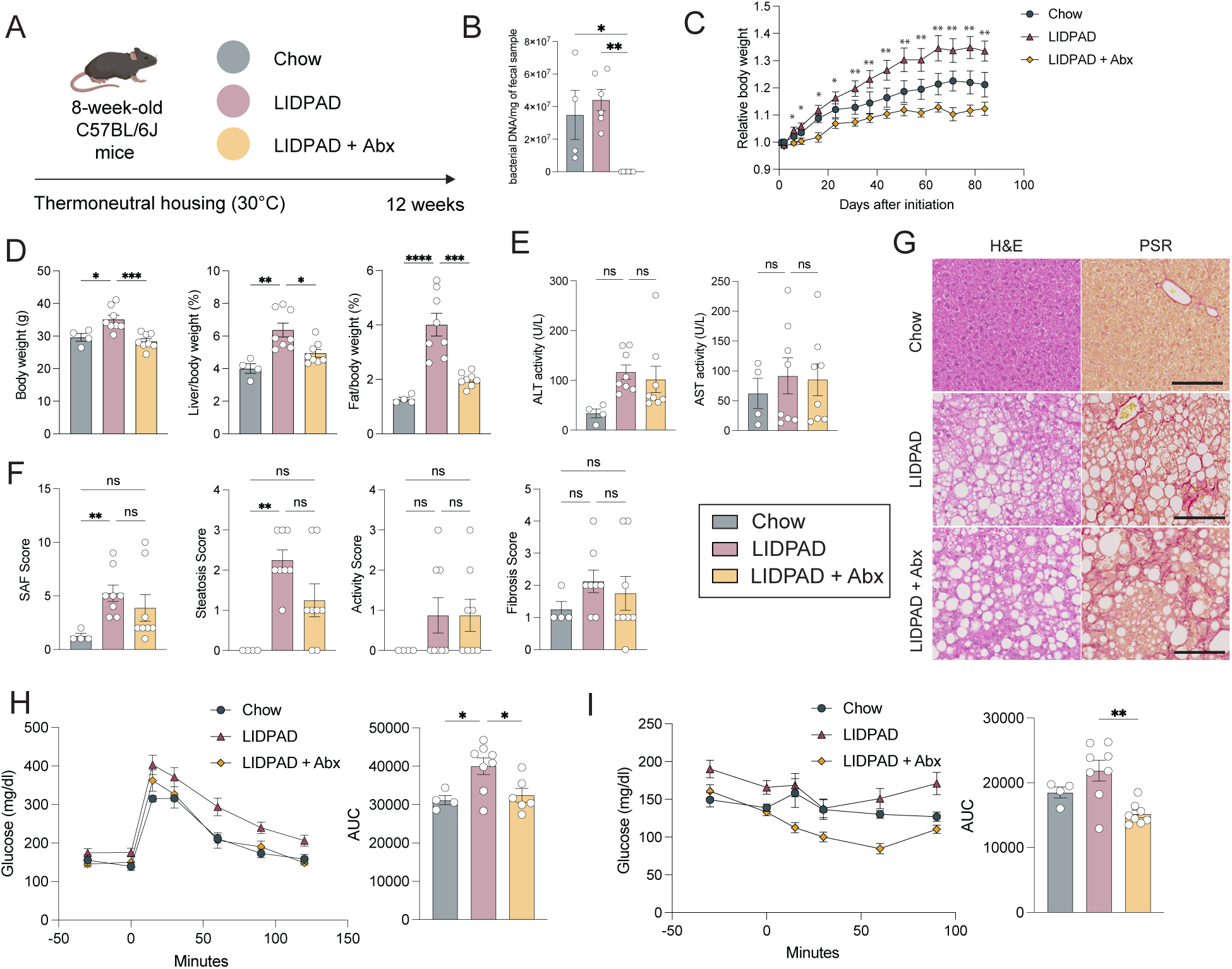
Gut microbiota depletion modulates metabolic phenotypes in the standard LIDPAD model. **(A)** Schematic of the experimental design. Eight-week-old C57BL/6J mice were fed chow, a LIDPAD diet, or a LIDPAD diet combined with a mixture of antibiotics (LIDPAD + Abx) for 12 weeks under thermoneutral housing conditions (30°C). **(B)** Faecal bacterial DNA content per mg of faecal sample, quantified by qPCR. **(C)** Body weight normalised to initial body weight over the study period. **(D)** Body weight, liver/body weight ratio, and fat/body weight ratio at the end of study. **(E)** Plasma ALT and AST activities. **(F)** Composite SAF (steatosis, activity, fibrosis) score, steatosis score, activity score and fibrosis score determined by histological assessment of liver sections. **(G)** Representative histological images of liver sections stained with haematoxylin and eosin (H&E) or picrosirius red (PSR). Scale bars, 50 μm. **(H)** Oral glucose tolerance test (OGTT) and corresponding area under the curve (AUC). **(I)** Insulin tolerance test (ITT) and corresponding AUC. Data are presented as mean ± SEM, with each point representing an individual mouse; n = 4 (chow), 8 (LIDPAD), 8 (LIDPAD + Abx). Statistical significance was determined by one-way ANOVA with Tukey’s post hoc test. *P < 0.05, **P < 0.01, ***P < 0.001, ****P < 0.0001; ns, not significant.

**Supplementary Figure 2.**
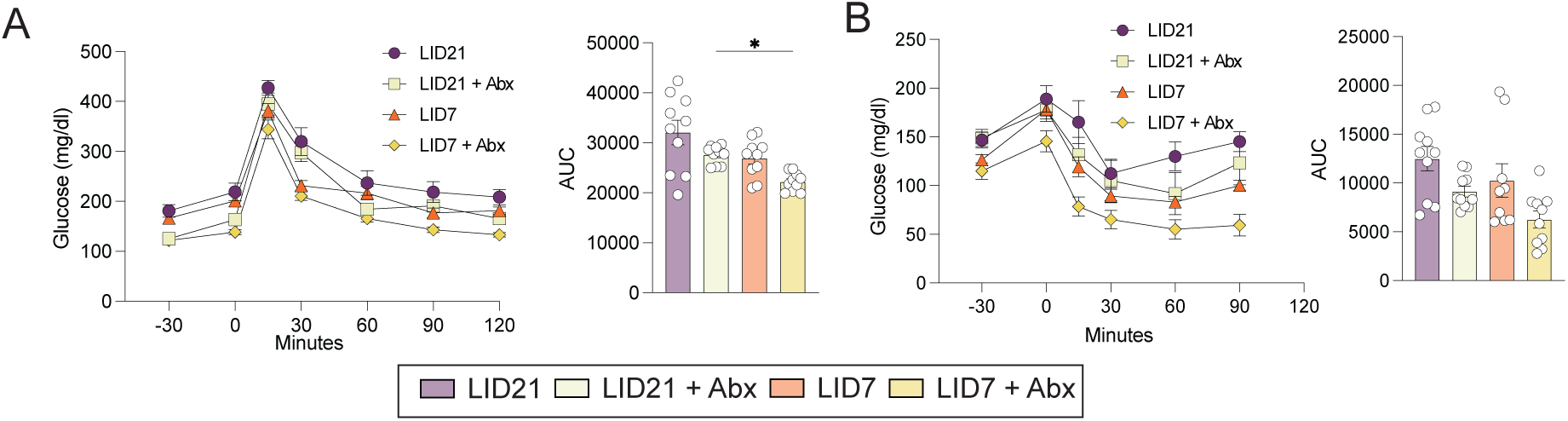
Glucose homeostasis in mice fed LID21 or LID7 diet with or without microbiota depletion. **(A)** Oral glucose tolerance test (OGTT) and corresponding area under the curve (AUC) in LID21, LID21 + Abx, LID7 and LID7 + Abx mice. **(B)** Insulin tolerance test (ITT) and corresponding AUC in the same groups. Related to Figure 1. Data are presented as mean ± SEM, with each point representing an individual mouse; n = 10 per group. Statistical significance was determined by one-way ANOVA with Tukey’s post hoc test. *P < 0.05; ns, not significant.

**Supplementary Figure 3.**
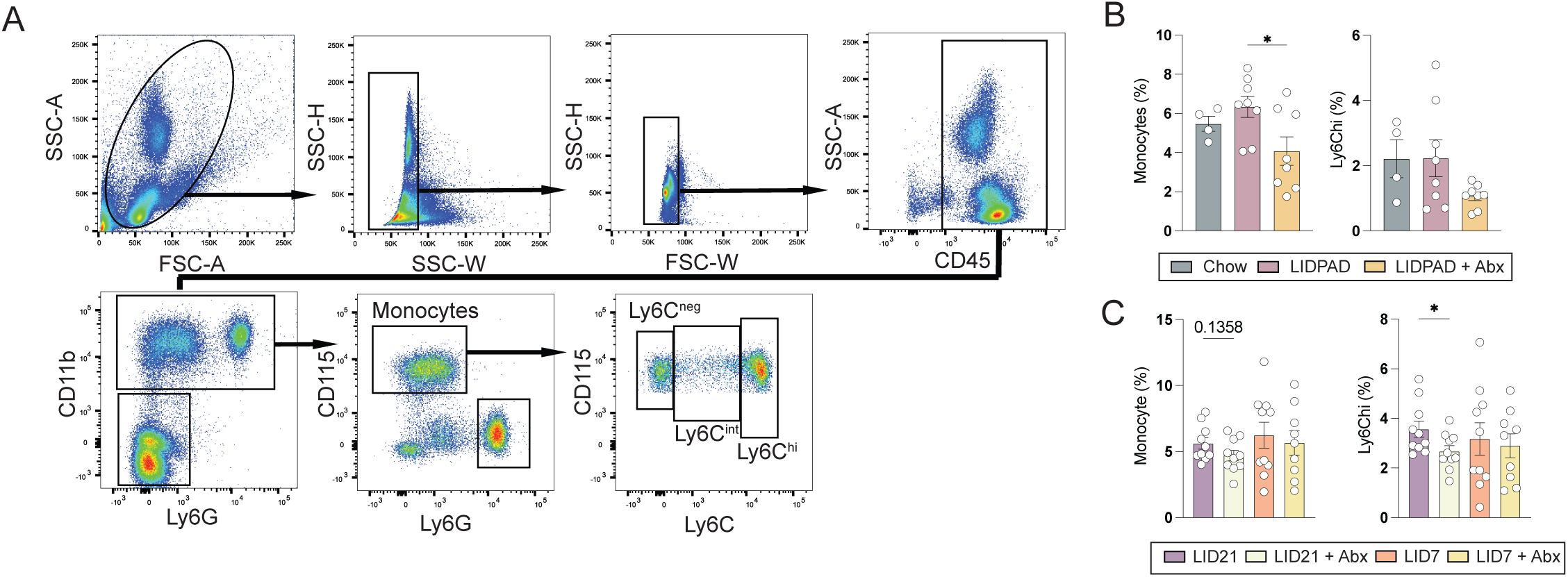
Circulating myeloid populations following microbiota depletion. **(A)** Representative gating strategy for flow cytometric identification of monocyte subsets in blood. Cells were gated on singlets, CD45⁺ leukocytes, CD11b⁺Ly6G⁻ myeloid cells and CD115⁺ monocytes, and monocyte subsets were resolved as Ly6C^high^, Ly6C^intermediate^ or Ly6C^negative^. **(B)** Frequencies of circulating monocytes and Ly6Cʰⁱ monocytes (% of CD45⁺ cells) in chow-, LIDPAD- and LIDPAD + Abx-fed mice. **(C)** Frequencies of circulating monocytes and Ly6Cʰⁱ monocytes (% of CD45⁺ cells) in LID21, LID21 + Abx, LID7 and LID7 + Abx mice. Data are presented as mean ± SEM, with each point representing an individual mouse; n = 4 (chow), 8 (LIDPAD), 8 (LIDPAD + Abx) for (B) and n = 10 per group for (C). Statistical significance was determined by one-way ANOVA with Tukey’s post hoc test. *P < 0.05; ns, not significant.

**Supplementary Figure 4.**
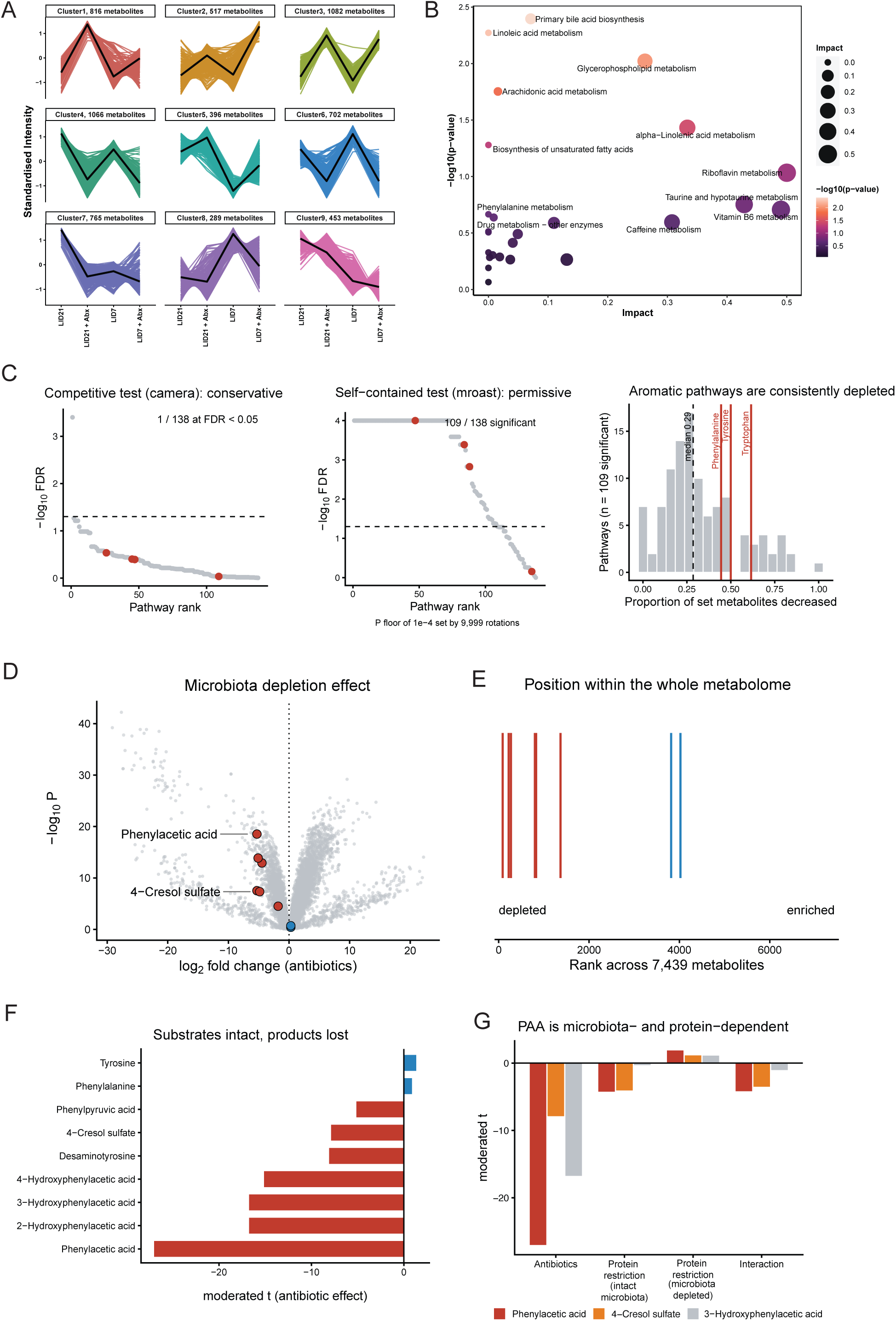
Caecal metabolite clustering and pathway enrichment across dietary and antibiotic groups. **(A)** k-means clustering of standardised caecal metabolite intensities across LID21, LID21 + Abx, LID7, and LID7 + Abx mice, yielding nine clusters based on shared abundance patterns across the four experimental groups. The number of metabolites assigned to each cluster is indicated. **(B)** Pathway enrichment analysis of the 1066 caecal metabolites assigned to cluster 4. Bubble size represents pathway impact, and colour represents statistical significance (−log10 p-value). **(C)** Pathway testing across all 138 KEGG pathways for the antibiotic contrast. Left, competitive test (camera), pathways ranked by significance. Middle, self-contained rotation test (mroast, 9,999 rotations); the plateau reflects the p-value floor set by the number of rotations. Dashed lines, FDR = 0.05. Red points, phenylalanine, tyrosine and tryptophan metabolism and their biosynthesis pathway. Right, distribution of the proportion of member metabolites decreased across the 109 pathways significant by mroast; dashed line, median; red lines, aromatic amino acid pathways. **(D)** Volcano plot of the antibiotic contrast across 7,439 metabolites (annotation levels 1 and 2, detected in ≥ 50% of samples). Red, microbial phenylalanine and tyrosine catabolites; blue, phenylalanine and tyrosine. Statistics are moderated t-tests from a two-factor limma model (∼ Protein × ABX). **(E)** All 7,439 metabolites ranked by moderated t statistic for the antibiotic contrast, from most depleted (left) to most enriched (right). Each vertical line marks one metabolite; colours as in (D). **(F)** Moderated t-statistics for the antibiotic contrast for individual aromatic amino acid metabolites. Colours as in (D). **(G)** Moderated t-statistics across the four model contrasts for phenylacetic acid, 4-cresol sulfate and 3-hydroxyphenylacetic acid.

**Supplementary Figure 5.**
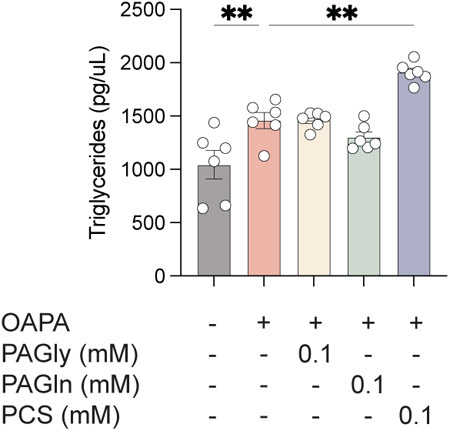
PAGln does not directly promote triglyceride accumulation in hepatocytes. Intracellular triglyceride content in hepatocytes treated with an oleic acid–palmitic acid mixture (OAPA) alone or in combination with PAGly, PAGln, or PCS (0.1 mM). Data are shown as mean ± SEM, with individual data points representing independent wells.

**Supplementary Figure 6.**
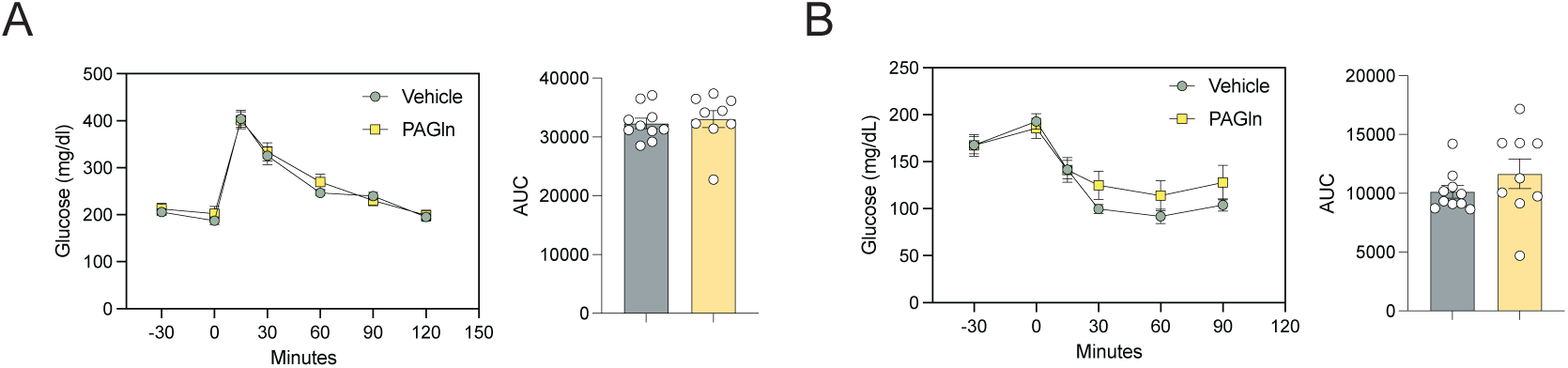
Glucose and insulin tolerance tests for supplementation of PAGln, related to Figure 3. **(A)** Oral glucose tolerance test (OGTT) and corresponding AUC in vehicle- and PAGln-treated mice. **(B)** Insulin tolerance test (ITT) and corresponding AUC in vehicle- and PAGln-treated mice. n = 10 (vehicle), 9 (PAGln). Statistical significance was determined by an unpaired two-tailed t-test.

**Supplementary Figure 7.**
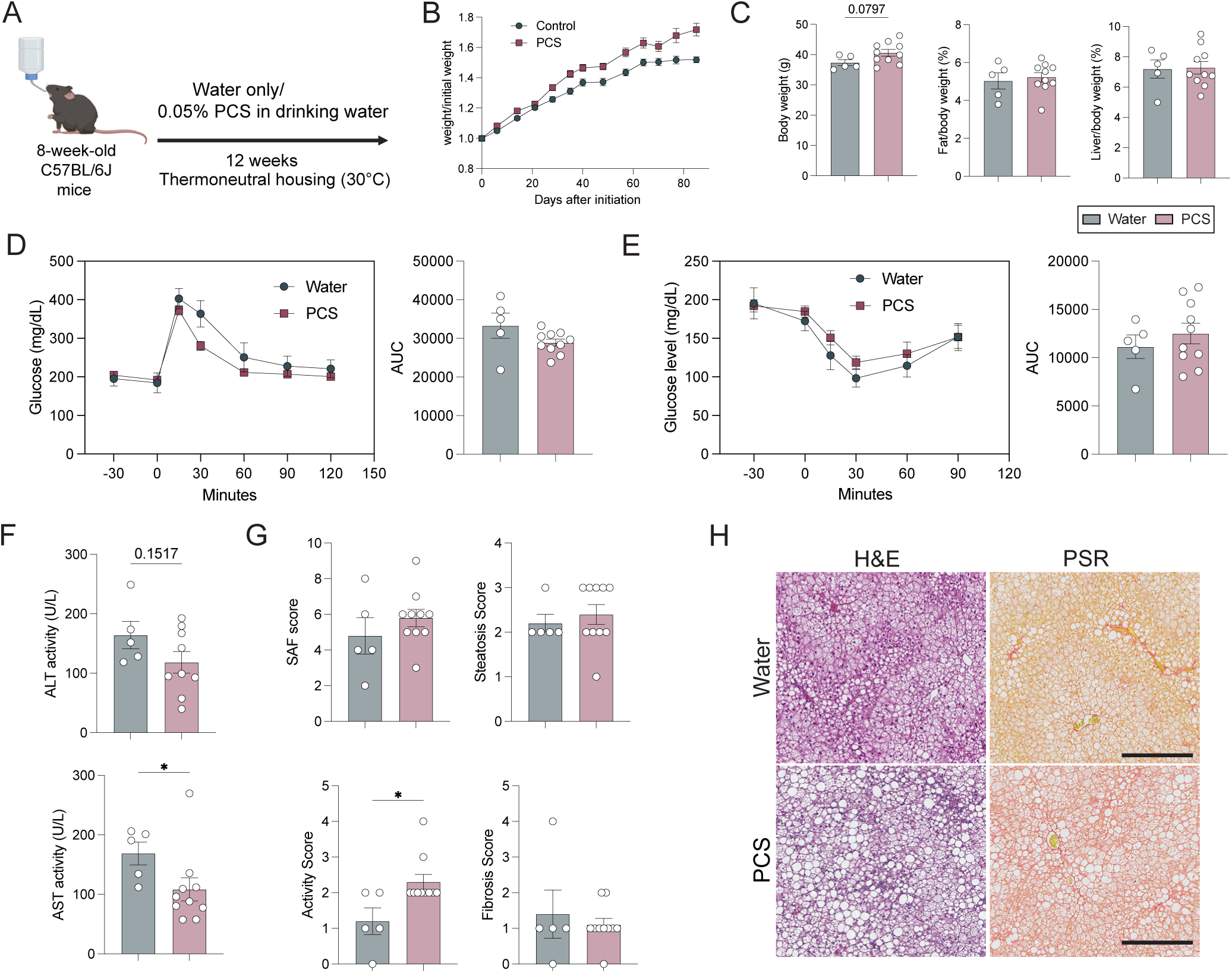
Oral PCS supplementation and metabolic phenotyping. **(A)** Schematic of the experimental design. Eight-week-old C57BL/6J male mice received water only or water supplemented with 0.05% p-cresol sulfate (PCS) while fed a LIDPAD diet under thermoneutral housing conditions for 12 weeks. **(B)** Body weight normalised to initial weight across the study period. **(C)** Final body weight, epididymal fat/body weight ratio and liver/body weight ratio in water- and PCS-treated mice. **(D)** OGTT with corresponding AUC in water- and PCS-treated mice. **(E)** ITT with corresponding AUC in water- and PCS-treated mice. **(F)** Plasma ALT and AST activities in water- and PCS-treated mice. **(G)** SAF score, steatosis score, activity score, and fibrosis score in water- and PCS-treated mice. **(H)** Representative histological images of liver sections stained with haematoxylin and eosin (H&E) and picrosirius red (PSR). Scale bars, 50 µm. Data are represented as mean ± SEM. n = 5 (water), 10 (PCS). Statistical significance was determined by unpaired two-tailed t test. *p < 0.05.

**Supplementary Figure 8.**
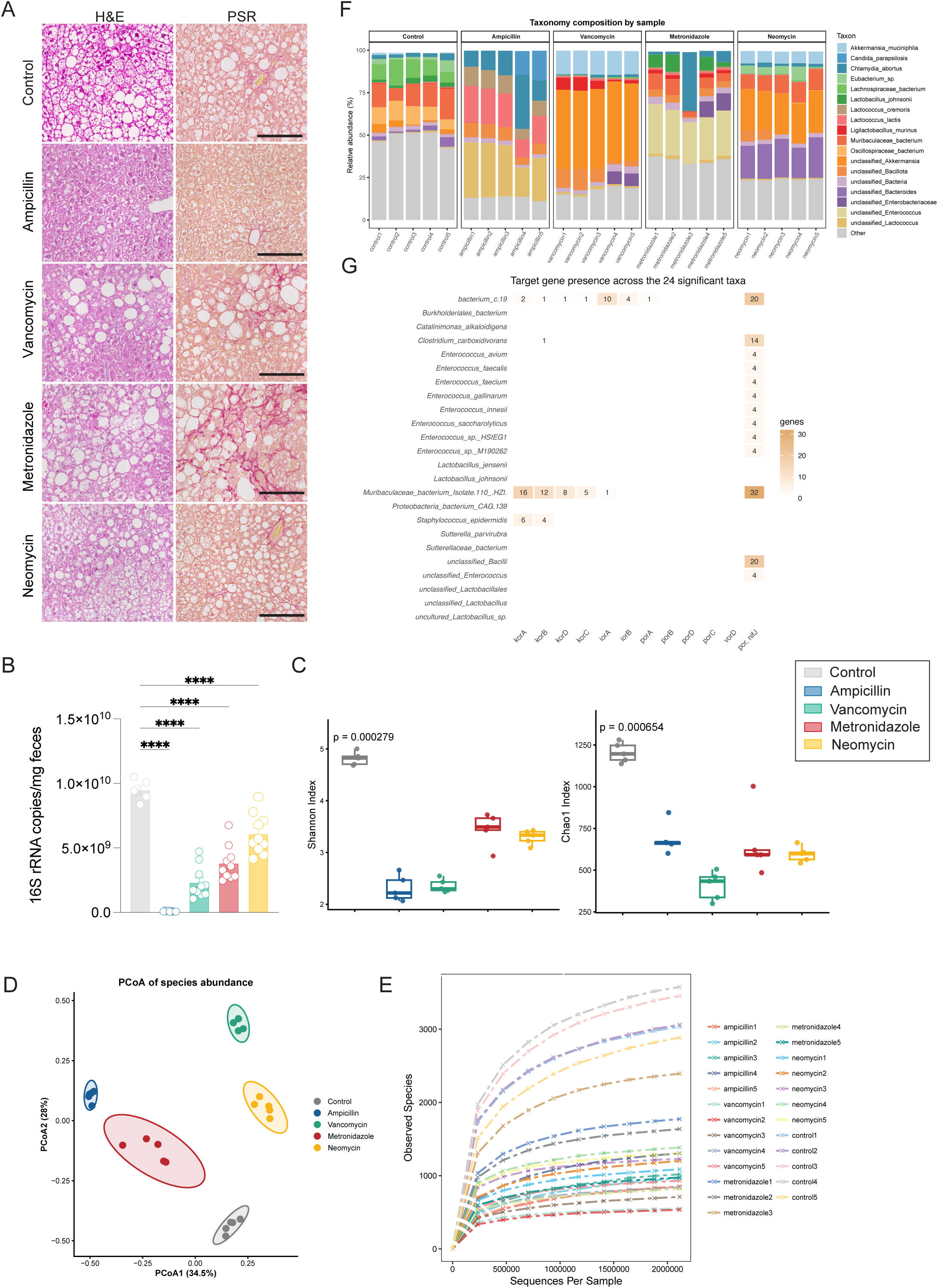
Individual antibiotics differentially alter gut microbiota composition and candidate phenylalanine catabolic gene content. **(A)** Representative haematoxylin and eosin (H&E) and picrosirius red (PSR) staining of liver sections from LID21-fed mice left untreated (Control) or treated with ampicillin, vancomycin, metronidazole, or neomycin. Scale bars, 50 μm. **(B)** Faecal bacterial 16S rRNA gene copy number (copies/mg faeces) quantified by qPCR across the five treatment groups. **(C)** Shannon and Chao1 diversity indices of the faecal microbiota across groups. **(D)** Principal coordinates analysis (PCoA) of faecal microbial community structure based on species abundance. **(E)** Species accumulation curves showing the number of observed species as a function of sequencing depth. **(F)** Relative abundance of bacterial taxa in individual faecal samples, grouped by treatment. **(G)** Heatmap showing the number of genes belonging to candidate phenylpyruvate/indolepyruvate:ferredoxin oxidoreductase gene families (*korA*-*D*, *iorA/B*, *porA*-*D*, *vorD*, *por*/*nifJ*) detected in the genomes of the 24 bacterial taxa significantly associated with both plasma ALT activity and SAF score. Cell colour intensity and overlaid numbers indicate gene counts. Data in (B) are presented as mean ± SEM, with each point representing an individual mouse. Statistical significance in (B) was determined by one-way ANOVA with Tukey’s post hoc test comparing antibiotic-treated groups with untreated controls (\*\*\*\**P* < 0.0001). *P* values in (C) are indicated on the plots.

**Supplementary Figure 9.**
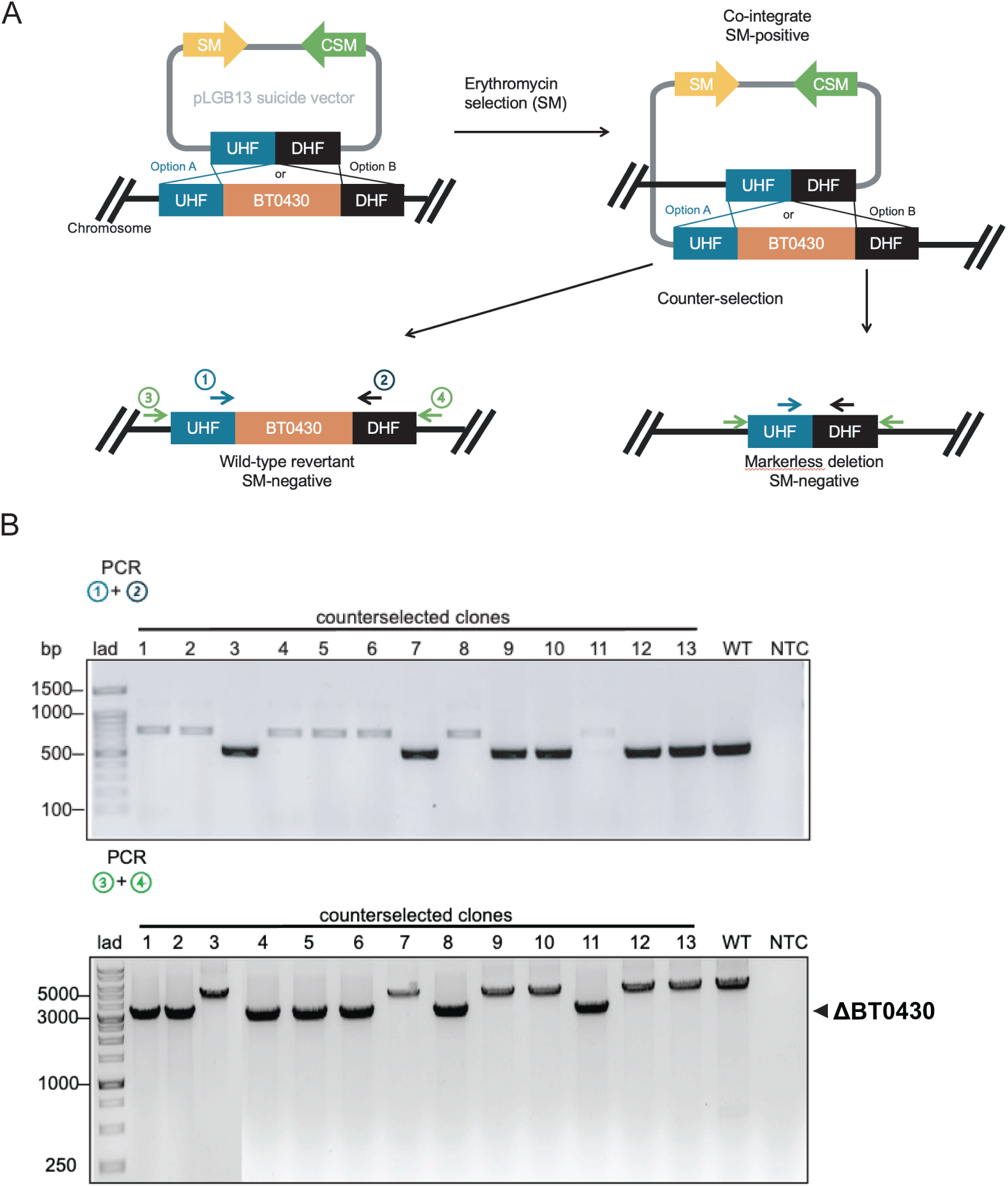
Generation and validation of the *B. thetaiotaomicron* ΔBT0430 mutant. **(A)** Schematic of the markerless allelic exchange strategy used to generate the ΔBT0430 mutant. The pLGB13 suicide vector carrying upstream (UHF) and downstream (DHF) homology flanks was integrated into the *BT0430* locus by homologous recombination following erythromycin selection (SM), generating a co-integrate. Counter-selection resolved the co-integrate into either a wild-type revertant or a markerless ΔBT0430 deletion. Numbered arrows indicate primer binding sites used for genotyping PCR. **(B)** Agarose gel electrophoresis of PCR products from counterselected clones using primer pairs 1+2 (top) and 3+4 (bottom), confirming the markerless ΔBT0430 deletion (arrowhead) relative to wild-type (WT) and no-template control (NTC).

## References

1. Younossi, Z. M. et al. Global epidemiology of nonalcoholic fatty liver disease-Meta-analytic assessment of prevalence, incidence, and outcomes. Hepatology 64, 73–84 (2016).

2. Kim, S. et al. Global burden of metabolic dysfunction-associated steatotic liver disease, 1990–2023, and projections to 2050: a systematic analysis for the Global Burden of Disease Study 2023. Lancet Gastroenterol. Hepatol. 11, 463–494 (2026).

3. Tacke, F. et al. EASL–EASD–EASO Clinical Practice Guidelines on the management of metabolic dysfunction-associated steatotic liver disease (MASLD). J. Hepatol. 81, 492–542 (2024).

4. Rinella, M. E. et al. AASLD Practice Guidance on the clinical assessment and management of nonalcoholic fatty liver disease. Hepatology 77, 1797–1835 (2023).

5. Byrne, C. D., Armandi, A., Pellegrinelli, V., Vidal-Puig, A. & Bugianesi, E. Μetabolic dysfunction-associated steatotic liver disease: a condition of heterogeneous metabolic risk factors, mechanisms and comorbidities requiring holistic treatment. Nat. Rev. Gastroenterol. Hepatol. 22, 314–328 (2025).

6. Huang, D. Q. et al. Type 2 diabetes, hepatic decompensation, and hepatocellular carcinoma in patients with non-alcoholic fatty liver disease: an individual participant-level data meta-analysis. Lancet Gastroenterol. Hepatol. 8, 829–836 (2023).

7. Choi, B. S.-Y. et al. Feeding diversified protein sources exacerbates hepatic insulin resistance via increased gut microbial branched-chain fatty acids and mTORC1 signaling in obese mice. Nat. Commun. 12, 3377 (2021).

8. Malik, V. S., Li, Y., Tobias, D. K., Pan, A. & Hu, F. B. Dietary Protein Intake and Risk of Type 2 Diabetes in US Men and Women. Am. J. Epidemiol. 183, 715–728 (2016).

9. Sluijs, I. et al. Dietary intake of total, animal, and vegetable protein and risk of type 2 diabetes in the European Prospective Investigation into Cancer and Nutrition (EPIC)-NL study. Diabetes Care 33, 43–48 (2010).

10. Newgard, C. B. Interplay between lipids and branched-chain amino acids in development of insulin resistance. Cell Metab. 15, 606–614 (2012).

11. Alferink, L. J. et al. Association of dietary macronutrient composition and non-alcoholic fatty liver disease in an ageing population: the Rotterdam Study. Gut 68, 1088–1098 (2019).

12. Lang, S. et al. High Protein Intake Is Associated With Histological Disease Activity in Patients With NAFLD. Hepatol. Commun. 4, 681–695 (2020).

13. Liao, Y. et al. Amino acid is a major carbon source for hepatic lipogenesis. Cell Metab. 36, 2437–2448.e8 (2024).

14. Hu, M., Xu, Y., Zhou, H. & He, X. Gut microbial metabolites of amino acids in liver diseases. Gut Microbes 17, 2586328 (2025).

15. Canfora, E. E., Meex, R. C. R., Venema, K. & Blaak, E. E. Gut microbial metabolites in obesity, NAFLD and T2DM. Nat. Rev. Endocrinol. 15, 261–273 (2019).

16. Tripathi, A. et al. The gut-liver axis and the intersection with the microbiome. Nat. Rev. Gastroenterol. Hepatol. 15, 397–411 (2018).

17. Nemet, I. et al. A Cardiovascular Disease-Linked Gut Microbial Metabolite Acts via Adrenergic Receptors. Cell 180, 862–877.e22 (2020).

18. Witkowski, M., Weeks, T. L. & Hazen, S. L. Gut Microbiota and Cardiovascular Disease. Circ. Res. 127, 553–570 (2020).

19. Zhu, Y. et al. Two distinct gut microbial pathways contribute to meta-organismal production of phenylacetylglutamine with links to cardiovascular disease. Cell Host Microbe 31, 18–32.e9 (2023).

20. Le Roy, T. et al. Intestinal microbiota determines development of non-alcoholic fatty liver disease in mice. Gut 62, 1787–1794 (2013).

21. Ishioka, M., Miura, K., Minami, S., Shimura, Y. & Ohnishi, H. Altered Gut Microbiota Composition and Immune Response in Experimental Steatohepatitis Mouse Models. Dig. Dis. Sci. 62, 396–406 (2017).

22. Hoyles, L. et al. Molecular phenomics and metagenomics of hepatic steatosis in non-diabetic obese women. Nat. Med. 24, 1070–1080 (2018).

23. Yang, M. et al. Western diet contributes to the pathogenesis of non-alcoholic steatohepatitis in male mice via remodeling gut microbiota and increasing production of 2-oleoylglycerol. Nat. Commun. 14, 228 (2023).

24. Parséus, A. et al. Microbiota-induced obesity requires farnesoid X receptor. Gut 66, 429–437 (2017).

25. Low, Z. S. et al. The LIDPAD Mouse Model Captures the Multisystem Interactions and Extrahepatic Complications in MASLD (Adv. Sci. 35/2024). Adv. Sci. 11, 2470214 (2024).

26. Yang, H. et al. Gut microbial-derived phenylacetylglutamine accelerates host cellular senescence. *Nat*. Aging 5, 401–418 (2025).

27. Romano, K. A. et al. Gut Microbiota-Generated Phenylacetylglutamine and Heart Failure. Circ. Heart Fail. 16, e009972 (2023).

28. Raverdy, V. et al. Data-driven cluster analysis identifies distinct types of metabolic dysfunction-associated steatotic liver disease. Nat. Med. 30, 3624–3633 (2024).

29. David, L. A. et al. Diet rapidly and reproducibly alters the human gut microbiome. Nature 505, 559–563 (2014).

30. Yao, C. K., Muir, J. G. & Gibson, P. R. Review article: insights into colonic protein fermentation, its modulation and potential health implications. Aliment. Pharmacol. Ther. 43, 181–196 (2016).

31. Diether, N. E. & Willing, B. P. Microbial Fermentation of Dietary Protein: An Important Factor in Diet^−^Microbe^−^Host Interaction. Microorganisms 7, 19 (2019).

32. Beaumont, M. et al. Quantity and source of dietary protein influence metabolite production by gut microbiota and rectal mucosa gene expression: a randomized, parallel, double-blind trial in overweight humans. Am. J. Clin. Nutr. 106, 1005–1019 (2017).

33. Henao-Mejia, J. et al. Inflammasome-mediated dysbiosis regulates progression of NAFLD and obesity. Nature 482, 179–185 (2012).

34. Long, Q. et al. Gut microbiota and metabolic biomarkers in metabolic dysfunction–associated steatotic liver disease. Hepatol. Commun. 8, e0310 (2024).

35. Rusman, R. D. et al. Gut microbiota and metabolic-associated steatosis liver disease: Unveiling mechanisms and opportunities for therapeutic intervention. World J. Exp. Med. 15, 107316 (2025).

36. Dodd, D. et al. A gut bacterial pathway metabolizes aromatic amino acids into nine circulating metabolites. Nature 551, 648–652 (2017).

37. Guo, C.-J. et al. Depletion of microbiome-derived molecules in the host using Clostridium genetics. Science 366, eaav1282 (2019).

38. Balsevich, G. et al. Stress-responsive FKBP51 regulates AKT2-AS160 signaling and metabolic function. Nat. Commun. 8, 1725 (2017).

39. García-Bayona, L. & Comstock, L. E. Streamlined Genetic Manipulation of Diverse Bacteroides and Parabacteroides Isolates from the Human Gut Microbiota. mBio 10, 10.1128/mbio.01762-19 (2019).

40. Han, J., Gagnon, S., Eckle, T. & Borchers, C. H. Metabolomic analysis of key central carbon metabolism carboxylic acids as their 3-nitrophenylhydrazones by UPLC/ESI-MS. Electrophoresis 34, 2891–2900 (2013).

41. Kreznar, J. H. et al. Host Genotype and Gut Microbiome Modulate Insulin Secretion and Diet-Induced Metabolic Phenotypes. Cell Rep. 18, 1739–1750 (2017).

42. Kanehisa, M. The KEGG database. Novartis Found. Symp. 247, 91–101; discussion 101-103, 119–128, 244–252 (2002).

